# The soil microbiome reduces Striga infection of sorghum by modulation of host-derived signaling molecules and root development

**DOI:** 10.1101/2022.11.06.515382

**Authors:** Dorota Kawa, Benjamin Thiombiano, Mahdere Shimels, Tamera Taylor, Aimee Walmsley, Hannah E Vahldick, Marcio FA Leite, Zayan Musa, Alexander Bucksch, Francisco Dini-Andreote, Alexander J Chen, Jiregna Daksa, Desalegn Etalo, Taye Tessema, Eiko E Kuramae, Jos M Raaijmakers, Harro Bouwmeester, Siobhan M Brady

## Abstract

*Sorghum bicolor* is one of the most important cereals in the world and a staple crop for smallholder famers in sub-Saharan Africa. However approximately 20% of sorghum yield is annually lost on the African continent due to infestation with the root parasitic weed *Striga hermonthica.* Existing Striga management strategies often show an inconsistent to low efficacy. Hence, novel and integrated approaches are needed as an alternative strategy. Here, we demonstrate that the soil microbiome suppresses Striga infection in sorghum. We associate this suppression with microbiome-mediated induction of root endodermal suberization and aerenchyma formation, and depletion of haustorium inducing factors (HIFs), root exudate compounds that are critical for the initial stages of Striga infection. We further identify microbial taxa associated with reduced Striga infection with concomitant changes in root cellular anatomy and differentiation as well as HIF degradation. Our study describes novel microbiome-mediated mechanisms of Striga suppression, encompassing repression of haustorium formation and induction of physical barriers in the host root tissue. These findings open new avenues to broaden the effectiveness of Striga management practices.

## Introduction

*Sorghum bicolor* is one of the most important cereal crops in the world as a source of food, feed, fiber and fuel. Its ability to withstand drought and soil aridity makes it a preferred crop in sub-Saharan Africa and earned it the name “the camel of crops” (Harris-Shultz et al., 2019). Despite its outstanding resilience to abiotic stresses, approximately 20% of sorghum yield is lost annually due to infestation with the root parasitic weed *Striga hermonthica* (Gurney et al., 1999). *Striga hermonthica* infects not only sorghum, but also many other crop species including rice, pearl millet and maize. An individual Striga plant can produce thousands of tiny, easy to spread seeds and its seedbank can remain dormant in soil for up to 20 years (Runo and Kuria, 2018). Striga is thus widespread in sub-Saharan Africa and its occurrence has been reported in at least 32 African countries (De Groote et al., 2008; Rodenburg et al., 2016). It is estimated that the annual cereal production losses amount to 6,213,000 tons of grain, worth $2.315 million USD annually (Rodenburg et al., 2016). These yield and economic losses often lead to field abandonment and food insecurity, which particularly affects smallholder farmers in sub-Saharan Africa.

The Striga life cycle is tightly connected to its host root chemistry. Upon phosphorus deprivation, host roots exude strigolactones, carotenoid-derived compounds that serve as a signal to recruit arbuscular mycorrhizal fungi. Striga has hijacked this strigolactone signal and germinates only upon its perception (Akiyama et al., 2005; Awad et al., 2006; Bouwmeester et al., 2020). Germinated Striga perceives other exudate compounds that act as haustorium inducing factors (HIFs, Bandaranayake et al., 2010). Haustorium development allows Striga to penetrate the host root tissue to reach its vasculature (Yoshida et al., 2016). Further establishment of a Striga xylem-host xylem connection is known as the “essence of the parasitism” (Kuijt, 1969). Through this xylem-xylem connection, Striga deprives its host plant from nutrients, water, and macromolecules, leading to adverse effects on plant growth and yield (Graves et al., 1989).

Currently, major practices of Striga management involve chemical control, “push-pull” methods, crop rotation and breeding for Striga-resistant host plant varieties. Despite these efforts, each management strategy has only partial Striga mitigation efficiency (Goldwasser and Rodenburg, 2013). Moreover, these measures are often not available to smallholder farmers in sub-Saharan countries where the most common solution is manual weed removal. Thus, there is a need for new and effective methods that can be integrated into current agricultural practices. Microbial-based solutions based on the soil suppressiveness phenomenon can meet these criteria.

Suppressiveness of soils to root diseases has been studied for bacterial, fungal and oomycete pathogens. In most cases the suppressiveness is microbial in nature as it can be eliminated by sterilization or pasteurization of the soil and can be transplanted to non-suppressive soils (Raaijmakers and Mazzola, 2016). In the disease-suppressive soils, despite the presence of a virulent pathogen, disease symptoms are less severe, do not occur at all, or the pathogen is able to initially cause a disease that later declines in severity (Weller et al., 2002). Mechanisms and causal microorganisms involved in disease suppressiveness to fungal root pathogens have been identified (Weller et al., 2002; Gomez Exposito et al., 2017). Little fundamental knowledge is available on the functional potential of the soil microbiome to interfere in the infection cycle of plant parasitic weeds.

Masteling et al. (2019) proposed several potential mechanisms by which microbes can suppress parasitic plant infection. Microbes can interfere directly with the parasite’s life cycle, by either their pathogenic effect on parasite seeds or by reduction of parasite seed germination and haustorium formation. The latter can occur via disruption of the biosynthesis or degradation of strigolactones and HIFs (Masteling et al., 2019). Microbes could also act indirectly, by affecting either the host plant itself or its environment. Microbes could enhance host nutrient acquisition and as a result reduce strigolactone exudation and, subsequently, parasite seed gemination. Alternatively, microbes could induce changes in root system or cellular architecture, providing an avoidance mechanism, or creating mechanical barriers, respectively. Lastly, microbes could also induce local or systemic resistance in the host plant (Masteling et al., 2019).

To date, several mechanisms by which microbes directly influence the *Striga* lifecycle have been described, including suppression of Striga seed germination by strains of *Pseudomonas* (Ahonsi et al., 2002), and infection of Striga by *Fusarium oxysporum f.sp. strigae* (Nzioki et al., 2016). Following these studies, a *Fusarium-based* inoculant has been developed and integrated into agricultural practices in Kenya, resulting in an increase in maize yield in Striga-infested fields (Nzioki et al., 2016). Thus far, indirect effects of the soil microbiome on Striga infection of sorghum have not been investigated and mechanistically resolved.

Here, we identify a soil whose microbiome reduces Striga infection in sorghum, and set out to describe its mechanism of action, with a focus on host root-related traits. We show that microbes in this soil degrade sorghum HIFs and subsequently hamper haustorium formation. Moreover, the soil microbiome induces changes in root cellular anatomy, including cortical aerenchyma formation and endodermal suberin deposition. We further identify specific microbial taxa within the soil and sorghum roots, that were associated with Striga suppression via these mechanisms. We validate these associations by testing individual bacterial isolates from the indicator taxa for their ability to degrade HIFs and induce structural changes in the host roots. Our data reveal that specific soil bacteria can induce multi-tiered protection against Striga and provide a foundation to harness the potential of microbes in an agricultural context.

## Results

### The soil microbiome impedes the post-germination stages of Striga infection

To explore the existence of Striga soil suppressiveness we selected a soil from the Netherlands, referred to as the “Clue Field” soil (Schlemper et al., 2017). This soil has previously been shown to differentially influence the rhizosphere community composition of two sorghum varieties with distinct susceptibility to *Striga hermonthica,* (Gobena et al., 2017; Schlemper et al., 2017; Kawa et al., 2021). We gamma-irradiated a batch of this soil for the purpose of sterilization and ensured that gamma sterilization did not affect the physico-chemical properties of the soil (**Supplementary Data 1**). We profiled the soil microbiome composition by sequencing 16S rRNA gene (bacteria) and ITS region (fungi) amplicons from the DNA extracted from bulk non-irradiated and gamma-irradiated soil. The alpha-diversity of the microbial composition of the non-irradiated soil was higher than that of the gamma-irradiated soil **(Fig. 1A),** while fungal composition was comparable between the two soils (**Fig. 1B**). The gamma-irradiated soil will herein be referred to as “sterilized” soil and the non-irradiated soil as “natural” soil.

**Figure 1.**
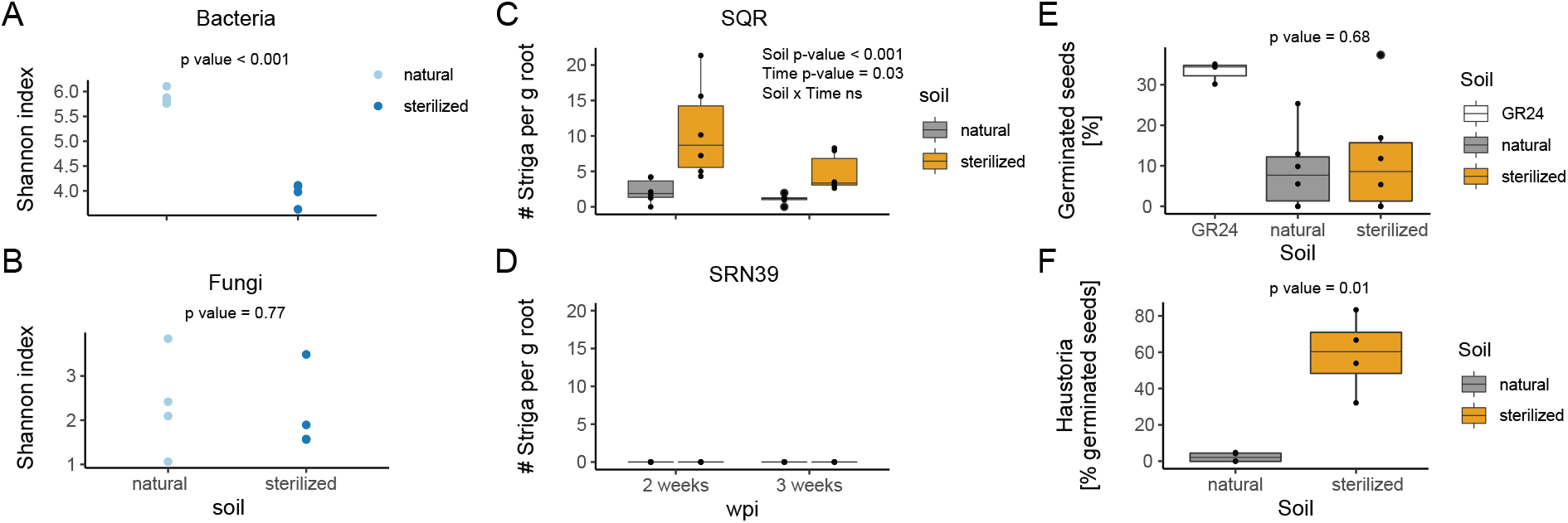
The soil microbiome suppresses Striga infection in sorghum. Alpha-diversity of (A) bacterial and (B) fungal communities of the field-collected soil (“natural”) and its gamma-irradiated counterpart (“sterilized”). Significance of the differences was determined with a Welch t-test (n=4). Number of Striga attachments per gram of fresh root weight of (C) Striga susceptible variety Shanqui Red (SQR) and (D) Striga resistant SRN39 at two and three weeks post-infection (wpi) in natural and sterilized soil. Significance of the differences was assessed with a two-way ANOVA (n=6). *In vitro* (E) germination and (F) haustorium formation of Striga seeds exposed to root exudates collected from four-week-old sorghum plants grown in natural and sterilized soil. The synthetic strigolactone, GR24, was used as a positive control for germination assay. Significance of the differences was assessed with a Welch t-test (exudates from six plants per soil were used with three technical replicates per exudate).

To test the effect of the soil microbiome on Striga infection in sorghum, we grew seedlings of Striga-susceptible Shanqui Red (SQR) and Striga-resistant SRN39 genotypes for ten days in 50 mL of either “natural” or “sterilized” soil to allow for microbial colonization of their roots (**Supplementary Fig. 1**). The 10-day-old seedlings, along with the soil “plug”, were transferred to larger pots with sand (control) or sand mixed with preconditioned *Striga hermonthica* seeds. The number of Striga attachments to sorghum roots were counted at two weeks and three weeks postinfection (wpi), which corresponds to four- and five-week-old plants, respectively. No Striga attachments were found on the roots of Striga-resistant SRN39 (**Fig. 1D**, **Supplementary Data 1**). We observed significantly fewer Striga attachments on SQR roots grown in the “natural” soil as opposed to the “sterilized” soil two and three weeks post-infection (**Fig. 1C, Supplementary Data 1**). This observation suggests that the Clue Field soil contains microbial components that partially suppress Striga infection.

Next, we asked at which stage of the Striga life cycle this suppression occurs. We set out to determine whether the functional outcome of the chemical signals governing Striga seed germination (host-derived strigolactones) and haustorium formation (host-derived HIFs) (Cui et al., 2018; Bouwmeester et al., 2020) is dependent on the soil microbial complement. To this end we collected root exudates from four-week-old SQR plants grown in the “natural” and “sterilized” soils in the absence of Striga and applied them to Striga seeds in an *in vitro* assay. We observed no difference in the germination percentage between seeds treated with sorghum root exudates from ‘‘natural” and “sterilized” soil (**Fig. 1E**). However, we noted a difference in the percentage of Striga seeds that formed haustoria. More than 60% of the Striga seeds exposed to the exudates from the “sterilized” soil developed haustoria whereas few haustoria were formed in the presence of the exudates of plants grown in the natural soil (**Fig. 1F**). Together, these *in vitro* results suggest that members of the “natural” soil microbiome reduce Striga infection of the susceptible sorghum cultivar at the post-germination stage of the parasite’s life cycle by interfering with haustorium initiation by the host-derived cues, the HIFs.

### The soil microbiome degrades haustorium inducing factors

The low level of haustorium induction by the root exudates from plants grown in the “natural” soil, suggests that the microbial component of this soil may influence the abundance of the HIFs. If this is the case, for translational purposes, ideally the microbes which reduce HIF levels should do so independent of Striga presence. To test this possibility, we measured the levels of known HIFs in the exudates of SQR plants grown in the “natural” and “sterilized” soil, both in the absence and presence of Striga. Using a two-way ANOVA, the differential abundance of detected HIFs and their dependence on the soil microbiome, Striga infection and their interaction, was determined. We detected five previously characterized HIFs with differential abundance in our treatments – acetosyringone, DMBQ (2,6-dimethoxybenzoquinone), syringic acid, vanillic acid and vanillin (Cui et al., 2018). Of these differentially abundant HIFs, syringic acid and vanillic acid levels were lower in exudates collected from “natural” soil compared to “sterilized” soil in the absence and presence of Striga at two weeks post-infection (**Fig. 2A, B**). Lower levels of DMBQ and acetosyringone were detected in the exudates of plants grown in “natural” soil, but only in the absence of Striga (**Fig. 2C-E**).

**Figure 2.**
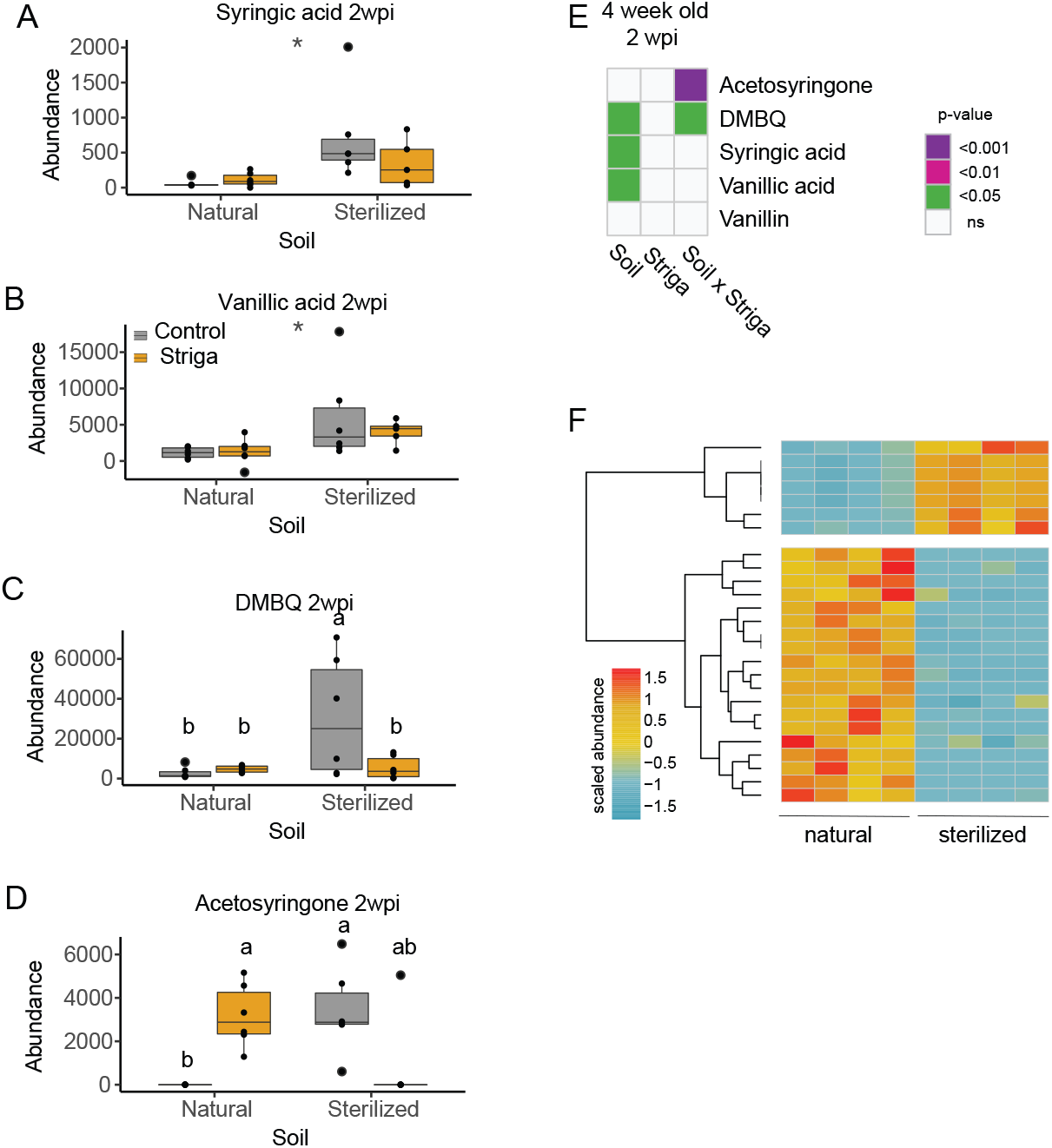
The soil microbiome influences haustorium inducing factor abudance in root exudates. Abundance of (A) syringic acid, (B) vanillic acid, (C) DMBQ, (D) acetosyringone at two weeks post-infection. Asterisks denote the significance of the soil impact, while different letters show significance of the differences between groups for traits, where soil-by-Striga interaction effect was detected (Tukey post-hoc test). E) Heatmap presenting the impact of the soil microbiome, Striga infection and their interaction on the abundance of haustorium inducing factors (HIFs) in root exudates as determined using a two-way ANOVA. Data presented are from two weeks post-infection with Striga, which corresponds to four-week-old sorghum plants. Purple, pink and green colors denote significant impact of the soil, Striga and their interaction. White squares indicate the lack of a significant effect (n=6). (F) Abundance of features identified with untargeted metabolite profiling, corresponding to potential HIF break-down products in root exudates collected from four-week-old plants grown in the “natural” and “sterilized” Clue Field soil (n=4). Values presented are the area under the associated peak scaled to the mean across all samples. The heatmap presents values for 26 compounds whose abundances differed significantly between exudates of plants grown in the two soils (p.adj <0.05, log_2_FC >1 or log_2_FC <-1).

Given the reduction of haustorium formation in exudates from “natural” soil in the *in vitro* assays (**Fig. 1F**) and the reduced levels of several HIFs in the exudates of plants two weeks post-infection (**Fig. 2A-D),** we hypothesized that microbes present in the “natural” soil degrade HIFs. To test this hypothesis, we used the BioTransformer database (Djoumbou-Feunang et al., 2019) to predict the products of potential microbial conversion of these HIFs (DMBQ, syringic acid, vanillic acid, vanillin, acetosyringone). In total 74 compounds were predicted as potential HIF break-down products (**Supplementary Data 1**). In the untargeted metabolite profiles of root exudates from four-week-old plants grown in “natural” or “sterilized” soil, we identified 82 features predicted to be HIF break-down products. Among these 82 features, abundances of 26 compounds differed significantly between exudates of plants grown in the “natural” or “sterilized” soils (p.adj <0.05, log_2_FC >1 or log_2_FC <-1). The majority (73%) of these compounds accumulated to higher levels in exudates from plants grown in “natural” than in “sterilized” soil (**Fig. 2F**). This indicates that in the presence of the soil microbiome from the “natural” soil, the putative HIF break-down products were more prevalent than the HIFs themselves. Collectively, we postulate that degradation of HIFs by members of the microbiome is associated with the reduction of Striga infection of sorghum plants grown in the “natural” soil.

### The soil microbiome modifies root cellular anatomy and corresponding transcriptional programs

Despite the ability of the soil microbiome to inhibit haustorium initiation, we still observed several Striga attachments on the roots of plants grown in “natural” soil (**Fig. 1C**). Thus, we next assessed whether the microbiome complement of this soil elicits additional changes in host root morphology that could influence Striga attachment and penetration. We conducted a detailed characterization of root system architecture and cellular anatomy to determine if any root traits are influenced by the soil microbiome, Striga infection or their interaction. Similar to HIF abundance in the root exudates, the soil microbiome affected the root traits in a manner independent of Striga infection (linear model term: soil) as well as in a more complex manner, dependently on Striga infection (linear model term: soil x Striga) (**Fig. 3A**). As with HIF abundances, we considered those traits that the microbiome changed independent of Striga for subsequent experiments.

**Figure 3.**
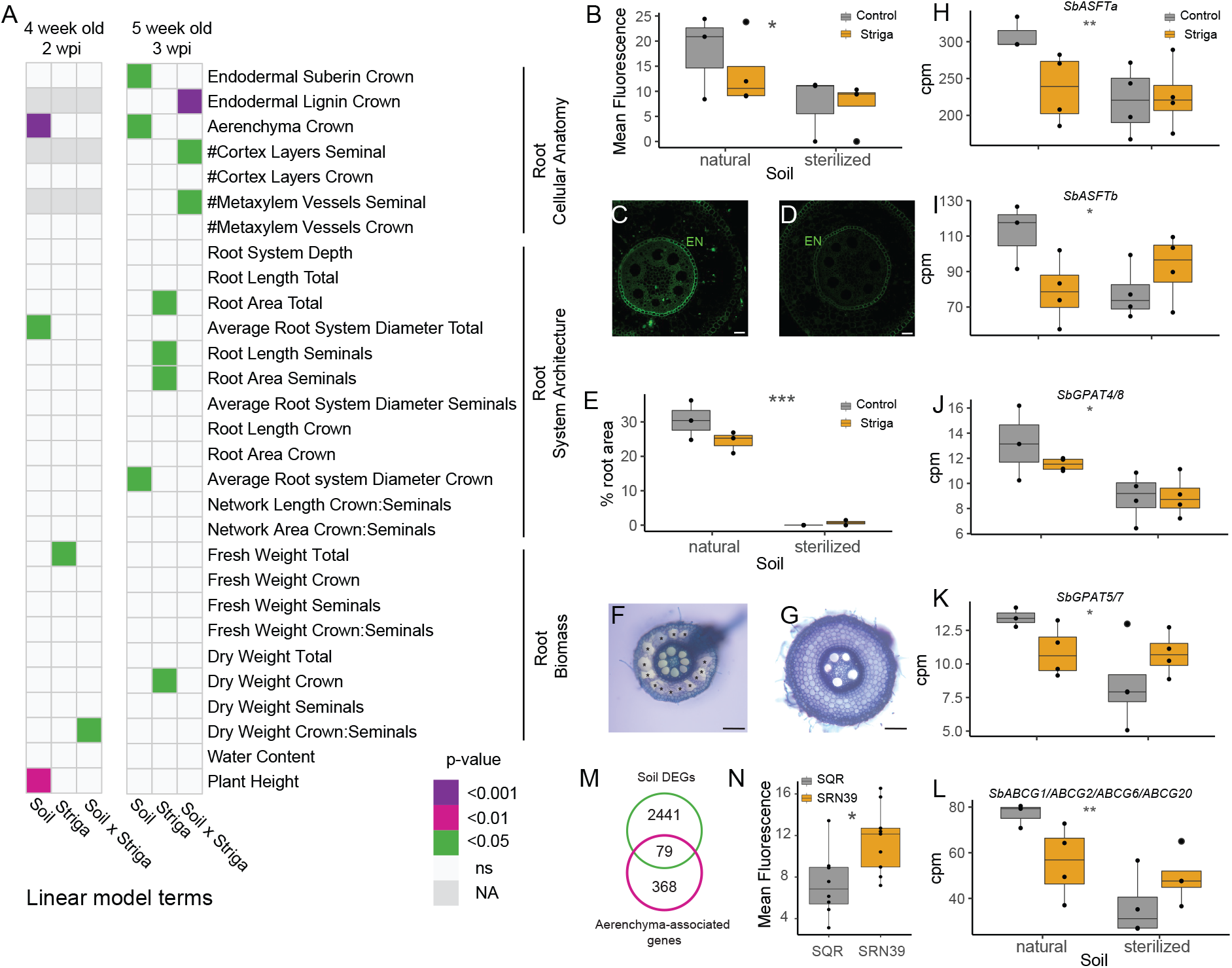
Striga suppressive soil induces changes in root system architecture and cellular anatomy. (A) Heatmap presenting the impact of the soil microbiome (Soil), Striga infection (Striga) and and their interaction (Soil x Striga) on root cellular anatomy, root system architecture and root biomass as determined by a two-way ANOVA. Data presented are from sorghum at two and three weeks post-infection (wpi) with Striga, which corresponds to four- and five-week-old sorghum plants. Purple, pink and green colors denote the significance threshold associated with the impact of the soil, Striga and their interaction on a given trait. White squares indicate a lack of significant effect, while NA denotes that trait was not tested at a given timepoint. Number or biological replicates tested for each trait is listed in **Supplementary Data 1**. (B, C, D) Suberin content in the endodermis of sorghum crown roots three weeks postinfection with Striga and (E, F, G) aerenchyma proportion in endodermis of sorghum crown roots three weeks post-infection with Striga grown on (C, F) natural and (D, G) sterilized soil. Suberin was stained with fluorol yellow and quantified by mean pixel fluorescence intensity. Aerenchyma area is expressed as a proportion of the whole root cross-section area. Asterisks in F indicate aerenchyma. Roots cross sections in F and G were stained with toluidine blue. Scale bar = 50 μm. Expression of suberin biosynthetic genes: (H) *SbASFTa,* (I) *SbASFTb,* (J) *SbGPAT4/8,* (K) *SbGPAT 5/7,* (L) *SbABSG1/ABSG2/ABCG6/ABCG20.* Expression of these genes was found to be regulated by the Clue Field soil microbiome (*adjusted p-value < 0.05, **adjusted p-value < 0.01). (M) Enrichment of sorghum orthologs of maize genes associated with root aerenchyma formation among the genes found to be regulated by the Clue Field soil microbiome (p-value = 0.008; Fisher’s exact test). (N) Endodermal suberization in roots of 10-day-old seedlings of SQR and SRN39, measured in 7 cm distance from the root tip (n = 12). The boxplots denote data spanning from the 25th to the 75th percentile and are centered to the data median. Dots represent individual values. Grey asterisks indicate the (B, E, L) p-value or adjusted p-value (H, I, J) for the term Genotype by a two-way ANOVA (B, E, L, H, I, J) or mixed model with experimental batch as a random factor (L). * p-value <0.05, ** p-value <0.01, *** p-value <0.001.

The root system architecture (RSA) of the mature sorghum plant consists of seminal and crown roots, thus each trait was quantified separately for crown and seminal roots, as well as for the entire root system (**Supplementary Fig. 1**). The soil microbiome had only a marginal effect on RSA and affected only the average diameter of the whole root system (at two weeks postinfection) or of the crown roots (at three weeks post-infection) (**Fig. 3A**). Although the goal of these experiments was to decipher the influence of the soil microbiome on Striga infection from the perspective of the host, a significant impact of Striga on RSA was observed. The effect of Striga on RSA was more pronounced at three weeks post-infection. Here, the total length and area of the root system and seminal root length were greater in plants infected with Striga when compared to non-infected plants, regardless of the soil type (**Fig. 3A, Supplementary Fig. 2 A, E, F**). We also observed a higher dry biomass of crown roots in plants three weeks post-infection, as compared to its non-infected control (**Fig. 3A, Supplementary Fig. 2 A, H**).

The soil microbiome primarily influenced root cellular anatomy traits (**Fig. 3A**). More endodermal suberization was observed in crown roots of five-week-old plants in “natural” soil independently from Striga infection (**Fig. 3B-D**). Additionally, more aerenchyma formed in crown roots of four- and five-week-old plants grown in “natural” soil, as compared to the “sterilized” soil independently from Striga infection status (**Fig. 3E-G**). Similar to acetosyringone and vanillin levels, several root anatomy traits varied in a complex way dependent on the interaction of the soil microbiome and on Striga infection, including the number of cortex layers and metaxylem vessels in seminal roots and lignification of the endodermis in crown roots (**Fig. 3A, Supplementary Fig. 2 A-D).**

Our data suggests that the microbial component of the “natural” soil promotes endodermal suberin deposition and aerenchyma formation, concomitant with suppression of Striga infection. To determine the potential molecular mechanisms that underly these changes, we conducted transcriptome profiling of the sorghum root systems at both two and three weeks post-infection and in the presence and absence of Striga. Indeed, the transcription of several sorghum orthologs of suberin biosynthetic genes or putative transporters - *Sobic.003G368100 (SbASFTa)* and *Sobic.005G122800 (SbASFTb)* (Gou et al., 2009; Molina et al., 2009), *Sobic.004G010300 (SbGPAT4/8)* (Canto-Pastor et al., 2022), *Sobic.009G16200 (SbGPAT5/7)* (Beisson et al., 2007), *Sobic.001G413700 (SbABCG1/SbABCG2/ABCG6/ABCG20a,* **Supplementary Fig. 3**) (Yadav et al., 2014) were upregulated in “natural” soil compared to “sterilized” soil in the presence and absence of Striga (**Fig. 3H-L**). Sorghum orthologs of maize genes previously reported as associated with aerenchyma formation, were also enriched among the genes upregulated in the “natural” versus “sterilized” soil (p-value = 0.008 per Fisher’s exact test, **Fig. 3M**). These transcriptome data align with the observed microbiome-mediated changes of host root cellular anatomy which in turn is correlated with reduced Striga infection.

### Microbial-mediated root cellular anatomy perturbation mimics differences observed in a sorghum resistant genotype

We previously found increased expression of genes associated with fatty acid biosynthesis and a higher abundance of suberin monomers and poly-hydroxy fatty acids in roots of SRN39 as compared to SQR (Kawa et al., 2021). Additionally, polymorphisms in genes related to suberin and wax-ester biosynthesis were previously found in sorghum varieties originating from regions of different levels of Striga infestation (Bellis et al., 2020). This led to the hypothesis that the distinct cell anatomy observed in the “natural” soil with reduced Striga infection may mimic cell anatomy observations found in the Striga resistant genotype, SRN39. Indeed, more suberin was deposited in the endodermis of SRN39 roots as compared to SQR (**Fig. 3N**). Aerenchyma content did not differ significantly between roots of SRN39 and SQR (**Supplementary Fig. 4**). This proves that at least some of the properties of root cellular anatomy associated with lower Striga infection in Striga resistant genotypes can be also induced by microbes.

### Identification of soil microbial taxa associated with Striga suppression

To identify microbial taxa associated with reduced Striga infection observed in the “natural” soil, we amplicon-sequenced the bacterial communities from: i) the bulk soil; ii) sorghum rhizosphere (soil directly surrounding the root system), (iii) roots growing in the soil plug (referred to as “soil plug-associated roots”), and (iv) roots growing into the sand (referred to as “sand-associated roots”) with or without Striga **(Supplementary Fig. 1).** We used covariance in bacterial taxa abundance across conditions (“natural” and “sterilized” soil in combination with presence and absence of Striga, across two times of infection) to determine their potential link with the Striga infection suppression via the three identified mechanisms in a generalized joint attribute modeling (GJAM) approach. The outputs of these models were mined to identify taxa whose relative abundance was negatively correlated with the number of Striga attachments and either negatively correlated with abundance of HIFs with reduced levels in the “natural” soil compared to the “sterilized” soil (vanillic acid, syringic acid) or positively correlated with suberin levels in the endodermis and aerenchyma proportion (**Supplementary Data 3**).

We first asked in which microbial sub-category the taxa predicted to induce each of these host root-related traits reside. We thus identified the most associated taxa (by the magnitude of residual correlation) for a given trait that are present in at least one microbial sub-category (see Methods). The majority of the top 100 bacterial taxa predicted to reduce Striga infection were found in the rhizosphere three weeks post-infection (**Supplementary Fig. 5 A**). The top 100 bacterial taxa associated with a reduction in HIF levels were found in both the rhizosphere and the soil plug-associated roots, while those predicted to induce aerenchyma formation and suberization resided in the soil plug-associated roots (**Supplementary Fig. 5 B-F**). No unique bacterial classes were linked to each of the studied traits. For each of the five traits (Striga attachment, aerenchyma, suberin, syringic acid and vanillic acid), the top-ranking bacterial taxa belonged to the Gammaproteobacteria, Deltaproteobacteria and Alphaproteobacteria, Bacteroidia and Actinobacteria classes (**Supplementary Fig. 5**).

Given the variety of activities bacteria may possess, we next set out to identify the taxa that were linked to changes in root cellular anatomy or HIF degradation as well as to the reduction of Striga infection We thus created a combined rank (see Methods) that summarizes the potential of a given taxa to reduce Striga infection via one of the identified mechanisms (**Supplementary Fig. 6**). For each bulk soil/root-associated sub-category (described above), we identified taxa positively associated with root cellular anatomy trait (suberin, aerenchyma) and for which the same taxa were negatively associated with Striga infection. In the case of HIFs, the taxa would be negatively associated with syringic or vanillic acid levels and the same taxa would be negatively associated with Striga infection. The majority of putative Striga-suppressive bacteria, regardless of the trait to which they were associated, belonged to the Actinobacteria and Proteobacteria phyla (**Supplementary Fig. 7**)

Most bacterial taxa whose abundance positively correlated with suberin or aerenchyma content, negatively correlated with the number of Striga attachments; in other words, more bacteria-induced aerenchyma/suberin coincided with less Striga attachments (**Supplementary Fig. 6)**. Conversely, the majority of bacterial taxa negatively correlated with suberin and aerenchyma were positively correlated with Striga infection. The majority of bacterial taxa associated with an increased in aerenchyma formation at three weeks post-infection were also associated with suberin induction (**Supplementary Fig. 7 A-C)**. Bacterial taxa of interest for further studies with the purpose of reducing Striga infection may be those that are associated with multiple mechanisms (**Supplementary Fig. 7 A-B).**

### Specific bacterial isolates prevent haustoria formation and induce suberization

A collection of bacterial strains from 35 genera has previously been established for the Clue Field soil (Kurm et al., 2019). From this collection, we prioritized bacterial isolates that matched, based on their taxonomic delineation and 16S amplicon sequence similarities, with the candidates (based on the combined ranking) identified by GJAM analysis. More specifically, we selected four *Pseudomonas* (VK1987, VK2039, VK2050, VK2070) and four *Arthrobacter* isolates (VK1979, VK2073, VK2105, VK2106) to determine if these isolates were able to induce changes in the root-related traits associated with Striga suppression (HIF abundance, suberization, and aerenchyma content). We first tested three *Pseudomonas* strains that were associated with HIF degradation for their ability to affect haustoria formation (**Supplementary Data 6**). In the presence of *Pseudomonas* isolate VK1987 only 2% of germinated *Striga* seeds exposed to syringic acid developed haustoria as opposed to 70% haustorium induction elicited by a mock control (media with no isolate) (**Fig. 4A**). However, *Pseudomonas* isolate VK1987 did not reduce haustoria formation in the presence of vanillic acid **(Fig. 4B).** Isolate VK1987 is therefore able to reduce haustorium initiation specifically via the syringic acid HIF. The two remaining *Pseudomonas* isolates tested, VK2050 and VK2070, did not reduce haustorium induction in presence of either vanillic acid or syringic acid (**Fig. 4A-B**). *Pseudomonas*, together with *Arthrobacter*, were also predicted by GJAM to reduce *Striga* infection via induction of aerenchyma (**Supplementary Data 6**). None of the four *Pseudomonas* and none of the four *Arthrobacter* isolates we tested reproducibly induced aerenchyma formation (**Supplementary Fig. 8**).

**Figure 4.**
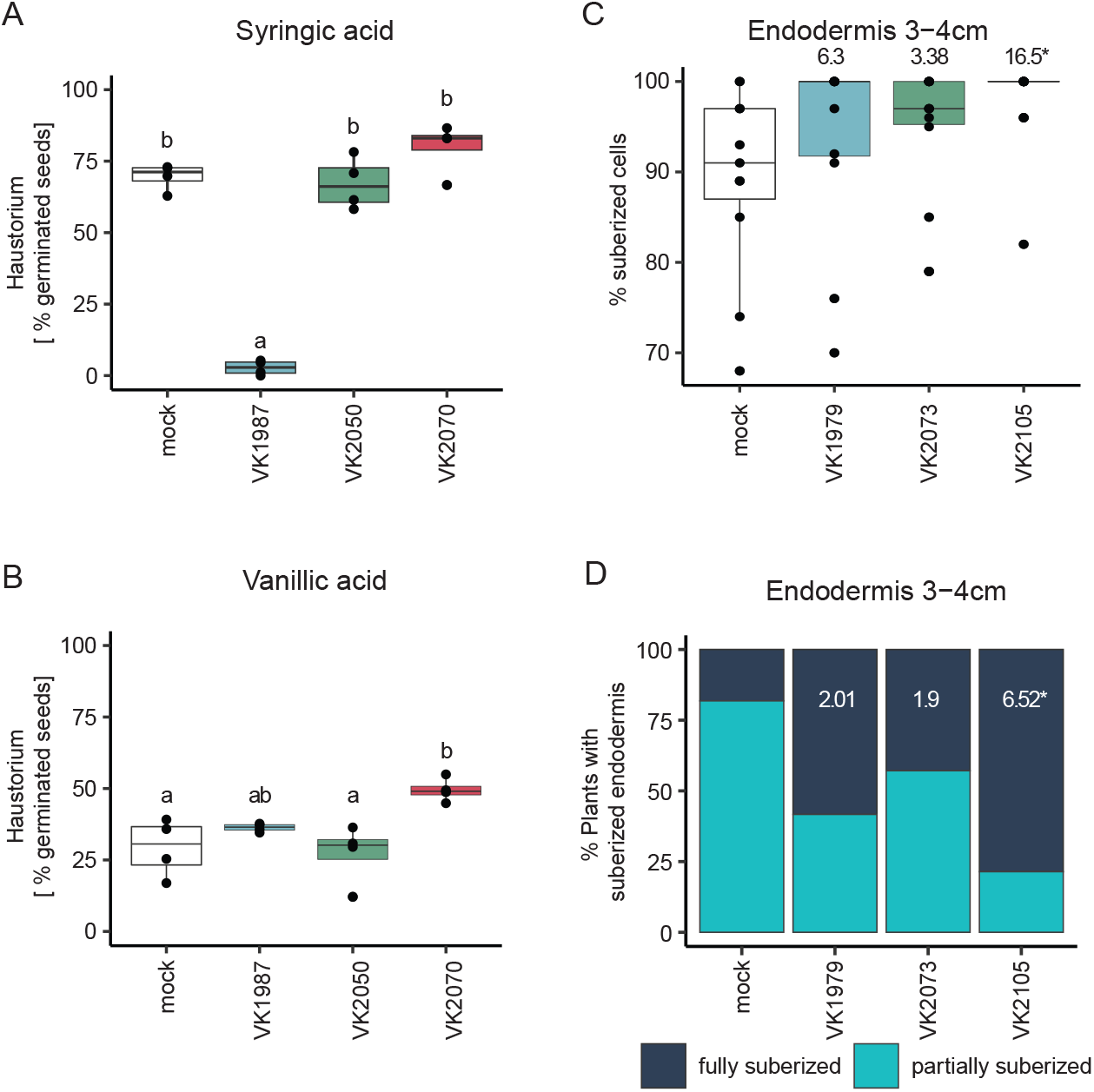
Individual bacterial isolates reduce haustorium formation and endodermal suberization. Percentage of germinated Striga seeds that developed haustorium in the presence of 100 μM (A) syringic acid and (B) vanillic acid incubated with *Pseudomonas* isolates VK1987, VK2050, VK2070. Sterile media used to grow the bacteria was used as a mock treatment. Statistical differences were tested with a one-way ANOVA and Tukey post-hoc test (p-value of the isolate effect: syringic acid <0.001, vanillic acid 0.0057). Letters denote significant differences between treatments. (C) Percentage of suberized cells in the endodermis and (D) percentage of plants with a fully and partially suberized endodermis within 3-4 cm from the root tip upon inoculation with *Arthrobacter* isolates VK1979, VK2073, VK2105. Numbers in C and D denote odds ratio and asterisks denote significant difference between plants inoculated with each isolate and the mock-treated plants determined by the least squares method.

*Arthrobacter* was the only genus associated (based on GJAM) with an increase in suberin content and Striga resistance, for which isolates were available in the Clue Field bacterial collection (**Supplementary Data 6**). In plant roots, suberin deposition occurs in three stages. In the first stage, there is an absence of suberin within the root meristem, followed by a “patchy” zone and a fully suberized zone in differentiated root (Kajala et al., 2021; Canto-Pastor et al., 2022). Typically, suberin in roots is quantified by measuring its level in few representative cells from the cross section (quantitative suberization), or by measuring the proportion of the non-suberized, “patchy” and fully suberized zone within the root length (developmental suberization) (Kajala et al., 2021; Canto-Pastor et al., 2022). The latter is challenging for sorghum, due to the high level of autofluorescence signal from its roots interfering with the suberin signal from fluorol yellow stain. We thus quantified the proportion of suberized cells within the endodermis in a radial crosssection and the proportion of plants with a fully suberized endodermis in a radial cross-section within the transition region between fully and “patchy” suberized zones (3-4 cm from the primary root tip, see Methods). We additionally quantified the effect of microbial inoculation on suberization of the exodermis, by quantifying the number of plants that developed a suberized exodermis.

Out of three tested *Pseudomonas* isolates (VK1979, VK2073, VK2105), none induced quantitative differences in the fully differentiated endodermis (6-7 cm from the root tip) and in the transition region between fully suberized and “patchy” zones (3-4 cm from the root tip, **Supplementary Fig. 9A**). However, in plants inoculated with *Arthrobacter* isolate VK2105 significantly more suberized cells were found within 3-4 cm from the root tip, thus in the region that constituted a “patchy” suberization zone in non-inoculated plants (**Fig. 4C**). Moreover, more plants developed a fully suberized endodermis when inoculated with *Arthrobacter* VK2105 isolate than in non-inoculated plants (**Fig. 4D**). Nearly 80% of the sorghum plants inoculated with *Arthrobacter* isolate VK2105 developed fully suberized endodermis, as opposed to 20% observed for the mock treatment (**Fig. 4D**). This increase in suberization extended to the root exodermis with a slightly precocious deposition of suberin in the exodermis (**Supplementary Fig. 9B**). Together, this demonstrates that individual bacterial strains are sufficient to perturb both the timing of suberin deposition as well as the number of suberized cells within the endodermal or exodermal cell files.

## Discussion

Here, we report that the microbiome of a field soil contributes to suppression of Striga infection of sorghum roots via disruption of host-parasite signaling and modulation of host root anatomy. Root exudates from sorghum grown in the Striga-suppressive soil did not affect Striga seed germination and strigolactone levels, but did significantly reduce haustorium formation, a phenotype that was associated with reduced levels of four key HIFs (syringic acid, vanillic acid, DMBQ, acetosyringone) (**Fig. 1, 2**). These results indicate that host-parasite signaling was disrupted at the level of haustoria formation via HIF degradation. Furthermore, more aerenchyma and endodermal suberin was detected in roots of sorghum grown in the presence of the soil microbiome (**Fig. 3**). These structural changes in root cellular anatomy likely affect the ability of Striga to penetrate the root. It is not known if progression of Striga through the root tissue requires a touch or mechanical stimulus from adjacent host tissue, but air-filled gaps in the cortex could likely disrupt this. Aerenchyma have also been associated with drought tolerance due to reduced metabolic and energy requirements (Zhu et al., 2010). An alternative hypothesis is that the parasitic plant may similarly sense a lack of metabolic activity and not continue with parasitization. A suberized endodermis can act as a physical barrier to Striga, preventing it from reaching the xylem. Physical barriers, consisting of lignin, callose, phenolic compounds or silica, can provide partial resistance in several host species and to several parasite species (Maiti et al., 1984; Rubiales et al., 2003; Gurney et al., 2006; Perez-de-Luque et al., 2006; Yoshida and Shirasu, 2009; Cissoko et al., 2011; Mbuvi et al., 2017; Mutinda et al., 2018; Mutuku et al., 2019). These barriers can be innate or induced upon infection with a parasitic plant (Kawa and Brady, 2022). Interestingly, accumulation of suberin was also observed for SRN39, a Striga-resistant sorghum genotype (**Fig. 3N**, Kawa et al., 2021). Whether this anatomical trait may be an additional postattachment resistance mode of other sorghum genotypes remains to be tested. Despite prior reports that root system architecture is associated with Striga resistance (Cherifari et al., 1990; Abate et al., 2017; Burridge et al., 2017), we observed no effect of the soil microbiome on RSA traits measured here (**Fig. 3A**).

To begin to validate the role of specific microbial taxa in HIF degradation and induction of suberization and aerenchyma, we tested a small number of bacterial isolates prioritized by GJAM analyses. It should be emphasized that the selection of the bacterial taxa was based on 16S amplicon sequence similarity, with only a minor 16S amplicon fragment as the template in the sequence alignment. In other words, our selection of the isolates was limited as it does not cover the full 16S sequence and, more importantly, does not use other taxonomic and functional markers of these bacterial taxa. Nevertheless, we did show that some of the bacterial isolates tested were able to degrade specific HIFs or to induce suberization (**Fig. 4**). These results validate, at least in part, that specific members of the soil microbiome can mediate post-germination (haustorium formation) and post-attachment (preventing Striga from penetrating through the root and thus establishing a vascular connection) resistance.

The potential of specific microbial species to degrade phenolic compounds, including vanillic and syringic acid has been previously reported (Wang et al., 2018; Oshlag et al., 2020). While it was mostly studied in the context of lignin decomposition, here we show that this activity can also be leveraged to reduce Striga parasitism. *Pseudomonas* isolate VK1987, that we associated with the reduction of HIFs in root exudates, inhibited haustorium formation in the presence of syringic, but not vanillic acid (**Fig. 4A-B**). Such selectivity towards some phenolic compounds among distinct microbes has been reported previously (Margesin et al., 2021).

Some pathogens reduce suberization of the endodermis that otherwise blocks their entry to plant vasculature (Froschel et al., 2021). Several commensal bacteria can lower suberin content in Arabidopsis (Salas-González et al., 2021). This negative effect of microbes on Arabidopsis suberization contrasts with our observations in sorghum, where multiple microbial taxa were associated positively with endodermal suberin content (**Supplementary Fig. 6**). These interspecies discrepancies could be caused by the presence of the exodermis in sorghum - an additional cell type where suberin can be deposited. Moreover, sorghum might assemble microbial communities different from those in Arabidopsis. Out of 41 endophytes tested in Arabidopsis, the majority reduced the fully suberized root zone, but six isolates promoted early endodermal suberization (Salas-González et al., 2021). This suggests that some overlap in suberin-inducing bacterial function might exist between sorghum and Arabidopsis.

Increased suberization could alternatively result from microbes affecting the nutritional status of the plant (indirect effect) or by microbes promoting the production of suberin precursors by the plant (direct effect). The latter has been observed in sorghum grown under drought (thus conditions promoting suberin deposition in roots (Baxter et al., 2009)), where increased production of glycerol-3-phosphate coincides with enrichment of monoderm bacteria, like Actinobacteria (Xu et al., 2018). It has been hypothesized that since monoderms use glycerol-3-phosphate to assemble their cell walls, they might induce its production in sorghum roots (Xu and Coleman-Derr, 2019). It is thus plausible that plants can also use this glycerol-3-phosphate as a substrate for suberin biosynthesis. Indeed, we observed Actinobacteria in our top ranked 100 taxa associated with suberization as well as upregulation of two glycerol phosphate transferases genes *(SbGPAT4/8* and *SbGPAT5/7)* in sorghum roots exposed to “natural” soil (**Fig. 3J, K**).

To the best of our knowledge, microbe-mediated induction of aerenchyma has not been reported. Ethylene induces aerenchyma formation (Yamauchi et al., 2013) and several ethylene-related genes were regulated by the Striga-suppressive soil microbiome (**Fig. 3M, Supplementary Data 2**). Microbes have been shown to interfere with plant ethylene signaling and both ethylene-producing and ethylene-degrading bacterial strains have been found (Ravanbakhsh et al., 2018). *A* reasonable hypothesis, therefore, is that ethylene-inducing microbes may induce aerenchyma that then restricts Striga entry into the root vasculature.

The known modes of pre- and post-attachment resistance in host species usually provide only partial protection. Likewise, each of the isolates identified her, on their own would likely provide little or limited protection against Striga. Combining microbes in a consortium that could induce multiple traits could provide a higher level of resistance. These microbial consortia should be assembled based on the extensive metagenomic sequencing and targeted identification and isolation of their members from the soils native to areas of their application. Bacterial taxa can then be used to prioritize the selection of microbial isolates from collections established from Striga-infested local soils in a targeted screens for resistance-associated phenotypes (HIF degradation, increase in aerenchyma and suberin content). Functional markers associated with i) the potential to degrade syringic acid, or (ii) upregulation of genes associated with the increase in aerenchyma content and suberization will further facilitate the targeted screens. While the host genotype-dependency of these identified mechanisms and their robustness to environmental conditions typical to areas where sorghum is grown still need to be addressed, this work lays the foundation for designing a multi-membered microbial consortium that suppresses haustorium formation and induces diverse structural barriers in roots to collectively reduce Striga parasitism.

## Materials and Methods

### Soil material

Soil was collected from the Clue Field in the Netherlands; 52° 03’ 37.91’’ N and 5° 45’ 7.074’’ E (Schlemper et al., 2017). Soil was dried and sieved through 4 mm mesh and one batch of it was sterilized by gamma irradiation with a dose of 8kGy, at room temperature by Steris (the Netherlands). Description of the physiochemical properties of “natural” and “sterilized” soils were provided by Eurofins Agro (the Netherlands, **Supplementary Data 1**).

### Plant material and growth conditions – soil “plug” assay

Seed of *Striga hermonthica* were collected in Sudan and kindly donated by Abdelgabar Babiker, Seeds were sieved by mesh of 200 μm pores to remove remaining soil particles and flower debris. Seeds were then surface sterilized with 10% (v/v) bleach and 0.02% (v/v) Tween-20 on filter paper and placed on a Buchner funnel connected to a vacuum pump until all liquid was removed. Next seeds were washed twice for 5 minutes in sterile water. Sterilized seeds were left to dry on the filter paper overnight in a laminar flow hood. Sterile seeds were mixed with sand containing around 16% (w/v) water and pre-conditioned for 10 days in a dark container in the greenhouse with temperature set to 26°C. As a negative control, sand without Striga seeds was treated in the same manner.

Seeds of *Sorghum bicolor* var. Shanqui Red (SQR) were obtained from GRIN (https://www.arsgrin.gov) and SRN39 seeds were kindly donated by Ethiopian Institute of Agricultural Research. Seeds were surface sterilized by agitating in a solution containing 4% (v/v) sodium hypochlorite and 0.2% (v/v) Tween-20 for 45 minutes followed by three rounds of 30 second incubations in 70% (v/v) ethanol followed by washes with sterile water. Next, seeds were washed four times with sterile water. Sterilized seeds were germinated on a wet Whatman paper (grade 1) at 28°C for 48 hours in the dark, followed by 48 hours in light. Four-day-old seedlings with approximately the same radicle length were transferred to 50 mL tubes filled with “natural” or “sterilized” soil (referred hereafter as the soil plug) mixed with 5% sterile water (w/v). Seedlings were watered with sterile water every second day. After 10 days, seedlings together with the soil plug were transferred to 40 cm long cones (Greenhouse Megastore, USA, catalog number CN-SS-DP) that were autoclaved prior to transfer. The bottom layer of the cones was filled with 350 mL of filter sand (0.5-1.0 mm, filcom.nl/) and the upper layer was filled with 350 mL preconditioned sand without (control) or with Striga seeds (3000 germinable Striga seeds per cone). Plants were organized in a randomized manner in the greenhouse compartment with the temperature set to 28°C during the day (11 hours) and 25°C at night (13 hours) with the 70% relative humidity and light intensity of 450μmol/m^2^/s. All measurements and sample collections were carried out at 14 and 21 days upon transfer to cones (referred to as two- or three-weeks post-infection; wpi). At day zero, seven and 14 (where day 0 is the day of the transfer to cones) plants were watered with 50 mL modified half-strength Hoagland solution containing 0.05 mM KH2PO4. On days one, four, 10, 13 and 17, plants were watered with 50 mL deionized sterile water.

### Striga infection quantification

Six individual plants of SQR and SRN39 were used per treatment (Striga-infected and control) at each time point (2,3 wpi). Sorghum plants were gently removed from the cones. All remaining sand and soil plug were collected and carefully examined for detached Striga plants. Roots were then gently washed in water and inspected under a dissecting microscope for early stages of Striga attachment. Roots were dried with a paper towel and fresh weight was recorded. The infection level was expressed as the ratio of total Striga attachments (the sum of early Striga attachments and the number of Striga plants recovered from the sand) and fresh root weight of individual plants. Data were analyzed with a two-way ANOVA, where a linear model was specified as: trait value=Soil+Treatment+Soil:Treatment, where Treatment stands for infected of not-infected (control) with Striga.

### Exudate collection and profiling

Each cone was flushed with water to collect 1 L of the flow-through. 100 mL of exudate were purified using solid phase extraction (SPE) with C18 Discovery® cartridges (bed wt. 500 mg, volume 6 mL, Merck). Cartridges were activated using 5 mL acetone and washed with 5 mL distilled water. 100 mL of sample were loaded on the cartridge and the flow through collected. The cartridge was further washed with 6 mL distilled water. Finally, compounds were eluted using 3 mL acetone. The acetone was evaporated using a SpeedVac (Scanvac, Labgene, Châtel-Saint-Denis, Switzerland). The semi-polar fraction of the exudates was reconstituted in 150 μL 25% (v/v) acetonitrile and filtered using a micropore filter (0.22 μm, 0.75 ml, Thermo scientific). The collected flow-through was freeze dried (Heto Powerdry LL1500, Thermo) and extracted with absolute methanol to remove the salts. The methanol was subsequently evaporated using a SpeedVac (Scanvac, Labgene, Châtel-Saint-Denis, Switzerland). The polar fraction of the exudates was reconstituted in 150 μL 25% (v/v) acetonitrile and filtered using a micropore filter (0.22 μm, 0.75 ml, Thermo scientific).

Untargeted analysis was performed as described in (Kawa et al., 2021). Briefly, 5 μL of root exudates (semi-polar and polar fraction) were injected on a Nexera UHPLC system (Shimadzu, Den Bosch, The Netherlands) coupled to a high-resolution quadrupole time-of-flight mass spectrometer (Q-TOF; maXis 4G, Bruker 194 Daltonics, Bruynvisweg 16 /18). Compounds were separated on a C18 stationary phase column. Peak finding, peak integration and retention time correction were performed as in (Kawa et al., 2021).

Targeted phenolics analysis was performed on a Waters Acquity UPLC™ I-Class System (Waters, Milford, MA, USA) equipped with Binary solvent manager and Sample manager was employed as a chromatographic system coupled to a Xevo® TQ-S tandem quadrupole mass spectrometer (Waters MS Technologies, Manchester, UK) with electrospray (ESI) ionization interface. five μL of root exudates (semi-polar and polar fraction) were separated on an Acquity UPLC™ BEH C18 column (2.1×100 mm, 1.7 μm particle size, Waters, Milford, MA, USA) with 15 mM formic acid in both water (A) and acetonitrile (B). At a flow rate of 300 μl per min and a column temperature of 40°C, the following gradient was applied: 0 min, 5% B; 2 min, 5% B; 32 min, 18% B; 60 min, 24% B; 65 min, 100% B. The compounds were measured in the ESI ion source of the tandem mass analyzer operating in the same conditions as in (Flokova et al., 2020). Mass data of phenolic compounds were acquired in multiple reaction monitoring (MRM) mode. The MassLynx™ software, version 4.1 (Waters), was used to control instrument and acquire and process MS data.

### Prediction of microbial degradation products

The structures of five HIFs: DMBQ, syringic acid, vanillic acid, vanillin, acetosyringone were inputted in the web-based tool BioTransformer to predict their microbial degradation products (http://biotransformer.ca/, (Djoumbou-Feunang et al., 2019). The first level predicted break-down compounds were used as an input for secondary break-down products. The exact masses of the degradation products were matched with the untargeted profiles of root exudates to retrieve potential candidates within a range on 25ppm error. Abundances of tested compounds in root exudates of plants grown in “natura” and “sterilized” soil (soil “plug” system) in the absence of Striga were compared with a Student’s t-test with false discovery rate adjustment from multiple comparisons.

### In vitro germination and haustorium formation assay

200 mg of Striga seeds were surface sterilized in 2% sodium hypochlorite containing 0.02% Tween-20 for 5 min, and then washed 5 times with sterile MilliQ water. The sterilized Striga seeds were spread on sterile glass fiber filter papers (Whatman ® GF/A, Sigma-Aldrich) in petri dishes moistened with 3 mL sterile MilliQ and preconditioned for 6-8 days at 30°C. 0.1 ppm GR24rac and 100 mM DMBQ was used as positive control for striga germination and haustorium formation, respectively. Stock of GR24rac was prepared in acetone and DMBQ was dissolved in methanol/water 50% (v/v). Dried, preconditioned Striga seeds were treated with 300 times-diluted root exudates from plants grown in the “natural” or “sterilized” soil, or GR24rac or DMBQ, and each of the solution was further divided into 3 technical replicates. Striga seeds were incubated in dark at 30°C for 2 days, when number of germinated Striga and haustoria formed were counted. The Striga germination rate was calculated for each replicate using the formula: GR% = (Ngs/Nts) × 100, where Ngs is the number of germinated seeds per well and Nts is the total number of seeds per well. The haustorium formation rate (HFR%) was calculated for each replicate using the formula: HFR % = (NHs/Ngs) × 100, where NHs is the total number of haustorium per well and Ngs is the number of germinated seeds per well. Welch t-sample test was used to compare effects elicited by the exudates of plants grown in the “natural” and “sterilized” soil.

### Root system architecture phenotyping

Data was collected 2 and 3 wpi, when plants were 4 and 5-week-old, respectively. Plant height was scored as a length from the sand surface to the bend of the highest leaf. Sorghum plants were removed from the cones and roots were cleaned from the sand and soil plug by gentle washes in water. Crown roots were separated from seminal roots and their fresh weight was scored separately. Roots were then placed in a transparent tray filled with water and scanned at 800dpi resolution with an Epson Perfection V700 scanner. Next, roots were dried with a paper towel, placed in paper bags, dried for 48 hours in 65°C and weighed to determine their dry weight.

Root system architecture was analyzed with the DIRT (Digital Imaging of Root Traits) software v1.1 (Das et al., 2015). The total root network area and total network length (to simplify we refer to it as total root area and total root length) used skeleton methods (Bucksch, 2014; Bucksch et al., 2014) as described in (Kawa et al., 2021). Mean root network diameter was calculated as the ratio of network area over network length. The dataset was cleaned from extreme outliers by removing individuals with values outside the 3^rd^ quartile. All collected data were analyzed with a two-way ANOVA, where a linear model was specified as: trait value=Soil+Treatment+Soil:Treatment, where Treatment stands for infected of not infected (control) with Striga.

### Root cellular anatomy phenotyping

Sorghum plants (2 and 3 wpi) were gently taken from the cones and washed in water to remove remaining sand and soil. For each plant a 1.5 cm segment of root tissue was cut from the tip of a crown root, from the middle of a crown root and from the middle of a seminal root. For the comparison of the root cellular anatomy of SRQ and SRN39 in a seedling stage, sterilized seeds were placed in 25 cm long germination pouches (PhytoAb Inc., catalog number: CYG-38LG) filled with 50 mL autoclaved water. Root tissue was harvested from 10-day-old seedlings. For each plant a segment of root tissue was cut from 7 cm distance from a root tip.

Root tissue was embedded in 5% (w/v) agar and fixed by a 10 minute vacuum infiltration in FAA solution (50% ethanol 95%, 5% glacial acetic acid, 10% formalin, 35% water, all v/v) followed by overnight incubation in FAA and rehydration by 30 minute incubations in a sequence of 70%, 50%, 30% and 10% (v/v) ethanol. Embedded tissue was stored in water at 4°C. Sections of 200-300 μm thickness were made with a Leica VT1000 vibrating microtome.

Suberin was stained with 0.01% (w/v) Fluorol Yellow 088 in lactic acid at room temperature, in the dark, for 30 min. Sections were rinsed three times for five minutes with water. Counter staining was done with 0.5% (w/v) aniline blue at room temperature for 30 minutes, followed by four 10-minute washes with water. Sections were mounted on slides with 50% glycerol prior to microscopic examination. Sections were imaged with LSM 700 laser scanning microscope (Carl Zeiss) with an excitation wavelength 488 nm and gain optimized to the signal strength. Quantification of endodermal suberin was done by calculating the mean fluorescence of two representative endodermal cells per section in ImageJ. Mean for two cells per section was used for further analysis.

For the aerenchyma quantification, separate set of sections were stained for 5 minutes in 0.1% toluidine blue (w/v) followed by five brief washes with water. Brightfield images were taken with Olympus AH-2. Aerenchyma proportion was expressed as the percentage of the area of the root section. The number of cortex layers and the number of metaxylem vessels were scored manually. The data collected from cones experiment were analyzed with a two-way ANOVA. For each genotype-time point data subsets a linear model was specified as: trait value=Soil+Treatment+Soil:Treatment, where Treatment stands for infected of not infected (control) with Striga. The data from the experiment with seedlings were derived from two independent experiments, thus a mixed model was used with experimental batch (Exp) as an independent factor specified with the formula: lmer (trait ~Genotype + (1|Exp)) with lme4 v.1.1-21 R package.

### RNA-seq library preparation

Two and three weeks after Striga infection (corresponding to 4- and 5-week-old plants) root material was harvested two hours after the light turned on. Each sorghum plant was gently removed from the cone and whole root system was cleaned from the remaining sand and soil by washing in water, dried with paper towel and snap frozen in liquid nitrogen (whole process took approximately 3 minutes per plant). Root tissue was ground with pestle in mortar, and RNA was extracted with RNaesy Plus Mini kit (Qiagen) with application of cell lysate on the QIAshredder columns (Qiagen) followed by the on-column Dnase I (Qiagen) treatment. Extracted RNA was precipitated with 3M NaOAc pH 5.2 (Thermo Scientific) in 100% ethanol and the pellet was washed with 70% (v/v) ethanol and dissolved in RNase-fee water. RNA-seq libraries were prepared with QuantSeq 3’ mRNA-Seq Library Prep Kit (Lexogen) following manufacturer protocol. Four biological replicates and three technical replicates for each RNA sample were used. Libraries were sequenced at the UC Davis DNA Technologies Core with Illumina HiSeq 4000 in SR100 mode.

### RNA-seq read processing and differential expression analysis

Quality control of obtained transcriptome sequences was determined with FastQC (http://www.bioinformat-ics.babraham.ac.uk/projects/fastqc/) before and after read processing. Three technical replicates of each library were pooled before reads processing. Barcodes were removed from raw reads with fastx-trimmer (http://hannonlab.cshl.edu/fastx_toolkit/index.html) with parameters: -v -f 12 -Q33. A reaper from Kraken Suite (Davis et al., 2013) was used for adaptor trimming and quality filtering with options: -geom no-bc -tabu $tabu -3pa $seqAdapt - noqc -dust-suffix 6/ACTG -dust-suffix-late 6/ACTG -nnn-check 1/1 -qqq-check 35/10 -clean-length 30 -polya 5. Processed reads were mapped to the reference genome of *Sorghum bicolor* BTx623 (McCormick et al., 2018) using STAR (Dobin et al., 2013) with options: -- outFilterMultimapNmax 20 --alignSJoverhangMin 8 --alignIntronMin 20 --alignIntronMax 10000 -- outFilterMismatchNmax 5 --outSAMtype BAM SortedByCoordinate --quantMode TranscriptomeSAM GeneCounts.

Genes for which no raw reads were detected across all samples were removed. Counts per million (CPM) were calculated with cpm() function from the edgeR package (Robinson et al., 2010). Only genes with a CPM > 1 in at least three samples were used for further analysis. CPM values are listed in **Supplementary Data 2**. Differentially expressed genes (DEGs) were determined with the R/Bioconductor limma package (Ritchie et al., 2015). CPM values were normalized with voomWithQualityWeights() function with quantile normalization to account for different RNA inputs and library sizes. Data from 4- and 5-week-old plants were analyzed separately. For each gene the linear model was defined as an interaction of the soil type (“natural” or “sterilized”) and treatment (control or infected with Striga) as: log(counts per million) of an individual gene ~ Soil*Treatment. Differentially expressed genes for each term of linear model were selected based on a false discovery rate < 0.05. Lists of differentially expressed genes for each term (soil, treatment, soil by treatment) are found in **Supplementary Data 2**.

### Gene orthology identification

List of sorghum orthologs of Arabidopsis suberin biosynthetic genes was obtained from (Canto-Pastor et al., 2022). To identify sorghum orthologs of ABCG transporter family proteins, a phylogenetic tree was generated as described in (Kajala et al., 2021). Next, we created a list of 672 maize genes whose expression was shown to change during root aerenchyma formation as reported in (Rajhi et al., 2011; Takahashi et al., 2015; Arora et al., 2017). Sorghum orthologs of maize genes were obtained from www.maizegdb.org. In total 447 unique sorghum genes have been defined as orthologs of maize genes associated with aerenchyma formation **Supplementary Data 2**. Enrichment of these genes among genes differentially expressed by soil type (2 wpi) was tested with Fisher’s Exact test.

### Microbial community analysis

The “natural” and “sterilized” soil plugs were prepared and placed in cones filled with sand like describe above, except no plant was transferred. The soil plugs and cones were placed in the same greenhouse compartment as cones with plants and were watered according to the same scheme. 14 days after the transfer to the cones, soil plug was excavated from the sand for the DNA extraction. These samples were used to profile microbiome communities of the bulk soil in the absence of plant.

The microbiome communities in the bulk soil in the presence of a plant and those associated with sorghum roots, bulk soil, rhizosphere and root material were collected 14 and 21 days after transfer to cones as in (Lundberg et al., 2012) with small modifications. First, soil not associated with roots was collected, shaken for 30 sec in 35 mL sterile phosphate buffer, centrifuged at 3,000 rpm for 20 min. Collected pellets were frozen in liquid nitrogen and constituted “bulk soil” samples. Roots that grew in the soil plug were carefully separated from roots that grew out of the soil plug and continued to grow in sand compartment. The excess of soil and sand was gently removed to leave a thin layer of 1-2 mm on the root surface. The roots were then shaken in 35 mL sterile phosphate buffer and transferred to a sterile petri dish containing phosphate buffer to be thoroughly washed and remove remaining soils/sand particles. The phosphate buffer was centrifuged at 3,000 rpm for 20 min, soil and sand pellets were frozen in liquid nitrogen and constitute a “rhizosphere soil” or “root-associated sand” samples. The fresh weight of washed roots was scored, and roots were immediately frozen in liquid nitrogen (and constituted “soil plug-associated roots” and “sand-associated roots” sub-categories).

The DNA was extracted with MoBio PowerSoil DNA isolation kit (Qiagen, Germany) from approximately 300 mg of grinded root material or 250 mg of soil/sand as recommended by the manufacturer. Prior to extraction from sand and soil, an additional centrifugation step was performed (10000 rpm at 4°C for 5 minutes). DNA concentrations were measured with a Nanodrop 2000 spectrophotometer (Thermo Fisher Scientific, USA) and stored at −80°C for further analysis. The DNA yield from “root-associated sand” was not sufficient for sequencing, thus these samples were discarded from further analysis.

Microbial communities were characterized by sequencing amplicons of the 16S rRNA region V3-V4 (with primer set: 16S_V3-341F: CCTACGGGNGGCWGCAG, 16S_V4-785R: GACTACHVGGGTATCTAATCC) for the bacteria, and ITS3-ITS4 (with primer set: ITS3_F: GCATCGATGAAGAACGCAGC, ITS4_R: TCCTCCGCTTATTGATATGC) for fungi. The amplicons were sequenced with Illumina MiSeq by BaseClear (Leiden, Netherlands). Raw sequence processing and quality control were performed with the UPARSE pipeline (Edgar, 2013). In brief, reads were paired and trimmed for quality (maximal expected errors of 0.25, reads length > 250 bp). Sequences were clustered into Operational Taxonomic Units (OTUs) at 97% of nucleotide identity, followed by chimera removal using UCHIME (Edgar et al., 2011). Taxonomic assignments of representative OTUs were obtained using the RDP classifier (Wang et al., 2007) against the Silva Database (Quast et al., 2013) (Quast et al., 2012). Sequences affiliated to chloroplasts were removed.

Analysis of microbial communities of “natural” and “sterilized” bulk soil without sorghum planted was performed with R phyloseq package v.1.26.1 (McMurdie and Holmes, 2013). Data was transformed with RLE normalization and rescaled to median sample count. Alpha diversity of each sample was calculated with estimate_richness function with “measures” set to “Shannon”. Significance of the difference between bulk “sterilized” and “natural” soil was determined with Student’s t-test.

### Identification of microbial candidates associated with reduced Striga infection

Statistical analyses were conducted in R v4.0.1 using different packages. To identify microbial taxonomic units associated with reduced Striga infection via identified phenotypes the generalized join attribute modeling was used with gjam package version2.6.2 (Clark et al., 2017) was used to estimate the effects of soil sterilization and Striga infection on the microbial communities (bacteria and fungi) within individual microbiome sub-categories (bulk soil, rhizosphere, soil plug-associated roots and sand-associated roots) and the number of Striga attachments and traits associated with Striga suppression (aerenchyma content, endodermal suberization, abundances of: syringic acid, vanillic acid and). The model analysis returns regression coefficients from the effect of the different treatments and quantified the increase or decrease in the microbial relative abundance and the changes in the other variables. Model diagnosis was evaluated the Markov Chain Monte Carlo (MCMC) to check when the estimated coefficients reached a stable value (after 10,000 simulations). Since the experiment consisted of a two-way factorial design, regression coefficients were compared against the following hypotheses: H1 - within each soil type (“natural” or “sterilized”) there is a difference between the Striga treatments (infected vs control); H2 - within each Striga treatments there is a difference between soil types. The model was applied for individual traits for the time point, at which they were found to be affected by the soil type, but not Striga infection, thus for 2 wpi: Striga attachments, aerenchyma content, abundances of: syringic acid, vanillic acid and; while for 3 wpi: Striga attachments, aerenchyma content, endodermal suberization).

As a joint model, gjam also allows to extract the residual correlations to investigate the relationship between the soil microbiome, Striga infection and associated traits (Leite and Kuramae, 2020). The residual correlations measure how strongly two different variables are associated regardless the influence of the treatment, which is therefore used to seek for potential biotic interactions (Pollock et al., 2014). In our study, residual correlations were calculated per each microbial subcategories and are listed in **Supplementary Data 3**.

Residual correlations calculated for individual microbial sub-category for each trait were filtered as follows: for number of Striga attachments and HIF abundances (vanillic acid, syringic acid) negative correlations, while for suberin content and aerenchyma proportion positive correlations were kept for further analysis. We first ranked the taxa based on their correlation for each trait across microbial sub-categories. Then, to identify taxa reducing Striga attachment number via each of identified mechanisms, ranking was done for each sub-category separately. Ranks for Striga attachment number and HIF levels were assigned so that the taxa with the lowest correlation received the highest rank value. Ranks for aerenchyma proportion and suberin content were assigned so that the taxa with the highest correlation received the highest rank value. Taxonomic membership was summarized for the top 100 bacterial taxa. The difference in the number of taxa that were summarized is due less fungal taxa were present in the Clue Field soil as compared to bacterial taxa.

Next a sum of ranks for Striga attachments number with rank for each one from: vanillic acid, syringic acid, suberin content, aerenchyma proportion, was calculated. The combined ranking was calculated as rank of this sum. Individual and listed ranks calculated per each microbial subcategory are presented in **Supplementary Data 4**. Taxonomic membership was summarized with a cut-off of residual correlation −0.2 for Striga attachments, syringic acid and vanillic acid levels and 0.2 for aerenchyma proportion and suberin content.

From the collection of bacterial strains isolated form the Clue Field soil by (Kurm et al., 2019) we selected isolates belonging to genera whose residual correlation were higher than 0.2 for suberin content and aerenchyma proportion and lower than −0.2 for Striga attachment number and HIFs abundances. By this we selected as candidates for reduction of Striga suppression via: i) promotion of aerenchyma formation: *Arthrobacter, Aeromicrobium, Bradyrizobium, Nocardia, Mesorhizobium, Paenibacillus, Phenylobacterium, Pseudomonas;* ii) degradation of vanillic and syringic acid: *Arthrobacter, Aeromicrobium, Pseudomonas* ii) induction of suberin deposition: *Arthrobacter*, *Aeromicrobium*, *Bradyrizobium*, *Nocardia*, *Mesorhizobium*, *Paenibacillus*. From the isolates we were able to re-grow and confirm their taxonomic identity by re-sequencing 16S rRNA (with primer set FQ 5’-AGAGTTTGATCCTGGCTCAG-3’ and REV 5’-GGTTACCTTGTTACGACTT-3’), four Pseudomonas, and four Arthrobacter isolates were used for in vitro tests of HIF degradation and inoculation of sorghum plants. Correlation residues derived from generalized join attribute modeling and ranks calculated for each genera in the collection can be found in **Supplementary Data** 5.

### Haustorium formation with individual bacterial isolates

The haustorium assay was conducted according to the protocol described by (Shimels et al., 2022). Briefly, a single bacterial colony was selected to inoculate minimal media Acetylglucosamine. The cultures were grown for 24 hrs at 25°C with shaking (200rpm). After adjusting the OD600 to 0.1, 10 μl of the overnight culture was added to 190ul of the acetyloglucosamine media supplemented with 100 μM of either syringic or vanillic acid (four biological replicates). After growth for another 24 hours, 50 μl of the cell-free culture filtrate was applied to pre-germinated Striga seeds to check the effect on haustorium formation. Striga seeds were prior surface sterilized with 0.5% sodium hypochlorite for 5 min and rinsed three times with sterile water. Approximately 100 seeds were then transferred to a wet 13 mm GF/A filter paper (VWR, Whatman). The seeds were then incubated at 30°C for 11 days in dark for preconditioning. To induce germination, 100 μl of water containing a final concentration of 1 μM GR24 was added. After 24 hours of incubation, pictures of the Striga seeds were taken and the percentage of seed that developed haustorium from all germinated seeds was scored using ImageJ. Four replicates were used per treatment. Statistical analyses were performed with a oneway ANOVA with a Tuckey post-hoc test.

### Sorghum inoculation with individual bacterial isolates

Individual isolates were cultured on a 1/10 dilution of tryptic soy broth (TSB, Difco) agar (1.5%, m/v) media containing 50 mg/L thiabendazole (Sigma) and incubated for 48 hours at 26°C. A single colony was then used to inoculate liquid TSB media (1/10 media dilution) and incubated for 48 hours at 26°C with shaking (200 rpm). Bacteria were harvested by centrifugation at 5000 rpm for 20 minutes and the resulting pellet was resuspended in sterile modified half-strength Hoagland solution (see methods for soil “plug” assay). In all the experiments individual isolates were applied to the sorghum variety Shanqui Red (SQR). An individual plant was inoculated with 10^7^ CFU/g sand in 5 mL of half-strength Hoagland media. The inoculum was applied at the root of a two-day-old sorghum seedling (pregerminated on wet Whatman paper for 48 hours at 28°C) at the same time as transplanting the seedling into a 50 mL tube filled with sand (moistened with 5 mL half-strength Hoagland media beforehand). Plants were watered every second day with 5 mL sterile water.

### Aerenchyma quantification in the presence of individual isolates

To estimate the proportion of aerenchyma we measured the porosity of the entire root system two weeks post inoculation following the protocol of (Van Noordwijk and Brouwer, 1988). Roots were gently removed from the sand, washed in water, very gently dried with a paper towel and weighed in 25 mL pycnometers (Eisco Labs) were filled with water and weighted. The harvested root systems were placed in individual pycnometer, refilled with water and weighted. Next, the pycnometers with roots were subjected to vacuum infiltration until the last air bubbles were seen, and their weight was scored. Root system porosity was calculated as: porosity = (P_v_ - P_r_)/ (P_w_+ RP_r_) where P_w_ is the weight of the pycnometer filled with water; P_r_ is the weight of the pycnometer filled with water and containing the root system; P_v_ is the weight of the pycnometer with a vacuum infiltrated root system and R is the root system weight at the moment of harvest. All tested isolates were first screened in two separate experiments with n= 6. Next, the isolates with largest different from the mock treatment were tested again with higher replication (n= 15).

### Suberin quantification – individual isolates assay

One week after bacterial inoculation roots were gently collected from the sand, washed in water and 1-1.5 cm of root segments were cut from two regions: 3-4 and 6-7cm from the root tip of the primary root. Region 3-4 cm constitutes the “patchy” suberization zone in mock-treated SQR root, while 6-7cm is the zone where the first onset of exodermis suberization is usually seen. Root tissue was embedded in agar, fixed in FAA, sectioned, and stained with fluorol yellow and imaged as described in the *Root cellular anatomy phenotyping* section. To quantify the differences along the root’s longitudinal axis, we also quantified the proportion of suberized and non-suberized cells in the endodermis. Given the technical challenges with obtaining sections that can be visualized in one plane, we excluded the areas of sections that were not completely perpendicular to the root’s longitudinal axis. These regions were determined by following the changes in the background fluorescent signal from the vasculature. The regions with less fluorescence in the vasculature, and adjacent endodermal cell were excluded from the analysis and are depicted in **Supplementary Fig. 10**. We also determined presence and absence of the suberized exodermis. Statistical analyses were done with a generalized linear model, for the proportion of plants with a fully suberized endodermis: glm(Fully_suberized ~ Strain, family = binomial(link = “logit”)), for the proportion of suberized cells: glm(cbind(‘Number of Suberized’,’Number of Non Suberized’) ~ Strain, family = quasibinomial(link = “logit”)), followed by comparison of each isolate with mock treatment with emmeans with option type = “response” with emmeans R package 1.8.1-1. The proportion of plants with a suberized exodermis was tested with Fisher’s exact test between each isolate and mock. Sections from 10-15 individual plants per treatment were used.

## Data availability

Sorghum sequences were deposited in NCBI GEO under the accession number GSE 216351.

Raw data from the growth measurements, root system architecture analysis, ANOVA tables and p-values for each statistical test can be found in **Supplementary Data 1**. CPM values and lists of differentially expressed genes are presented in **Supplementary Data 2.** Residual correlations from GJAM and ranks assigned to each microbial taxa are to be found in **Supplementary Data 3** and **Supplementary Data 4**, respectively. Residual correlations and rank for the members of the microbial collection are presented in **Supplementary Data 5**. Raw data and results of statistical analysis from the experiments with individual bacterial isolates can be found in **Supplementary Data 6**.

Data analysis scripts are publicly available at https://github.com/DorotaKawa/Striga-suppressivesoil. The script for generalized join attribute modeling can be found at https://github.com/Leitemfa/GJAM-PROMISE.

## Acknowledgments

DK, BT, MS, TT, AW, HEV, FD-A, JD, TT, EEK, JR, HB and SMB gratefully acknowledge support from the Bill and Melinda Gates Foundation, Seattle, WA via grant OPP1082853 “RSM Systems Biology for Sorghum”. SMB is partially funded by an HHMI Faculty Scholar grant and OS-1856749 and IOS-211980. AB was in part supported by the USDOE ARPA-E ROOTS Award Number DE-AR0000821 by the NSF CAREER Award No. 1845760 to A.B. Any Opinions, findings, and conclusions or recommendations expressed in this material are those of the author(s) and do not necessarily reflect those of the National Science Foundation. We would like to thank Ludek Tikovsky and Harold Lemereis for their assistance in the greenhouse, Angelica Guercio for help with quantification of root cellular anatomy and Daniel Runcie for the advice on the statistical analyses.

## Author Contributions

Conceptualization: DK, BT, MS, JMR, HB, SMB. Data curation: DK, BT. Formal analysis: DK, BT, MFAL, AB. Funding acquisition: HB, SMB. Investigation: DK, BT, MS, TT, AW, HEV, ZM, FD-A, AJC, JD, SMB. Methodology: DK, BT, MS, MFAL, AB. Project administration: EK, JMR, HB, SMB. Software: DK, MFAL, AB. Supervision: DK, BT, EEK, JMR, HB, SMB. Resources: TTe, JMR. Validation: DK, BT, MS. Visualization: DK. Writing – original draft: DK, SMB.

## Supplementary Figures

**Supplementary Figure 1.**
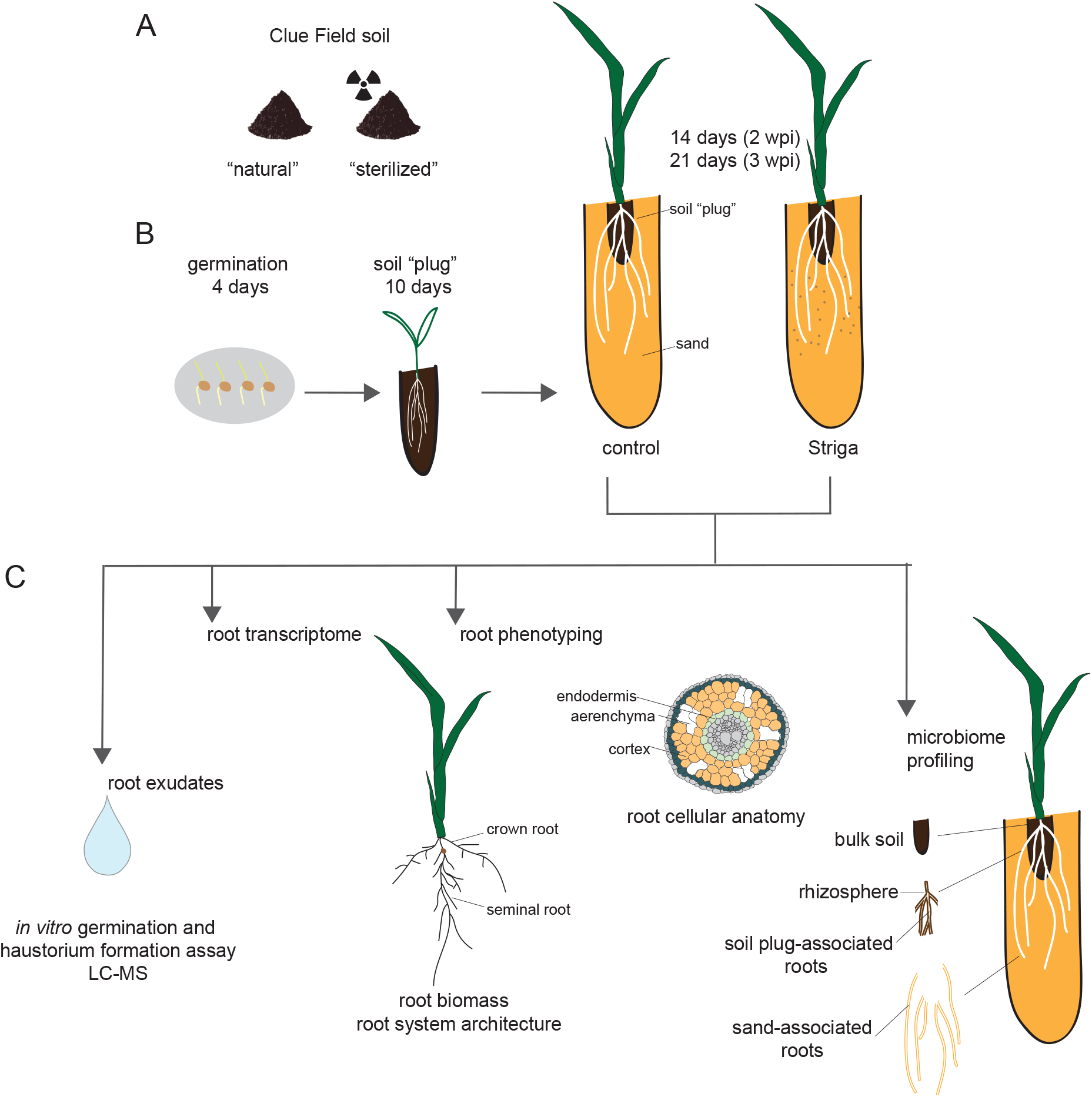
Visual description of the methodology used to investigate the mechanisms of Striga infection suppression by soil microbiome. (A) Two batches of Clue Field soil were tested: “natural” and “sterilized”, the latter subjected to gamma-irradiation. (B) Sorghum seedlings were germinated and grown for 4 days on moistened filter paper and then (B) transferred to a soil “plug” for 10 days. The seedling, together with the soil “plug” was then transferred to conical tubes filled with sand (control) or sand mixed with Striga seeds. (C) Root tissue and root exudates were collected two and three weeks post-infection (wpi). The root exudates were used for the *in vitro* germination and haustorium formation assay and metabolite analyses. Root phenotyping included quantification of root system architecture and cellular anatomy. Root transcriptomes were profiled with RNAseq. (D) Microbiome profiles were obtained from bulk soil, rhizosphere, soil plug-associated roots (root system part in contact with soil “plug”) and sand-associated roots (the root system that was in contact with sand).

**Supplementary Figure 2.**
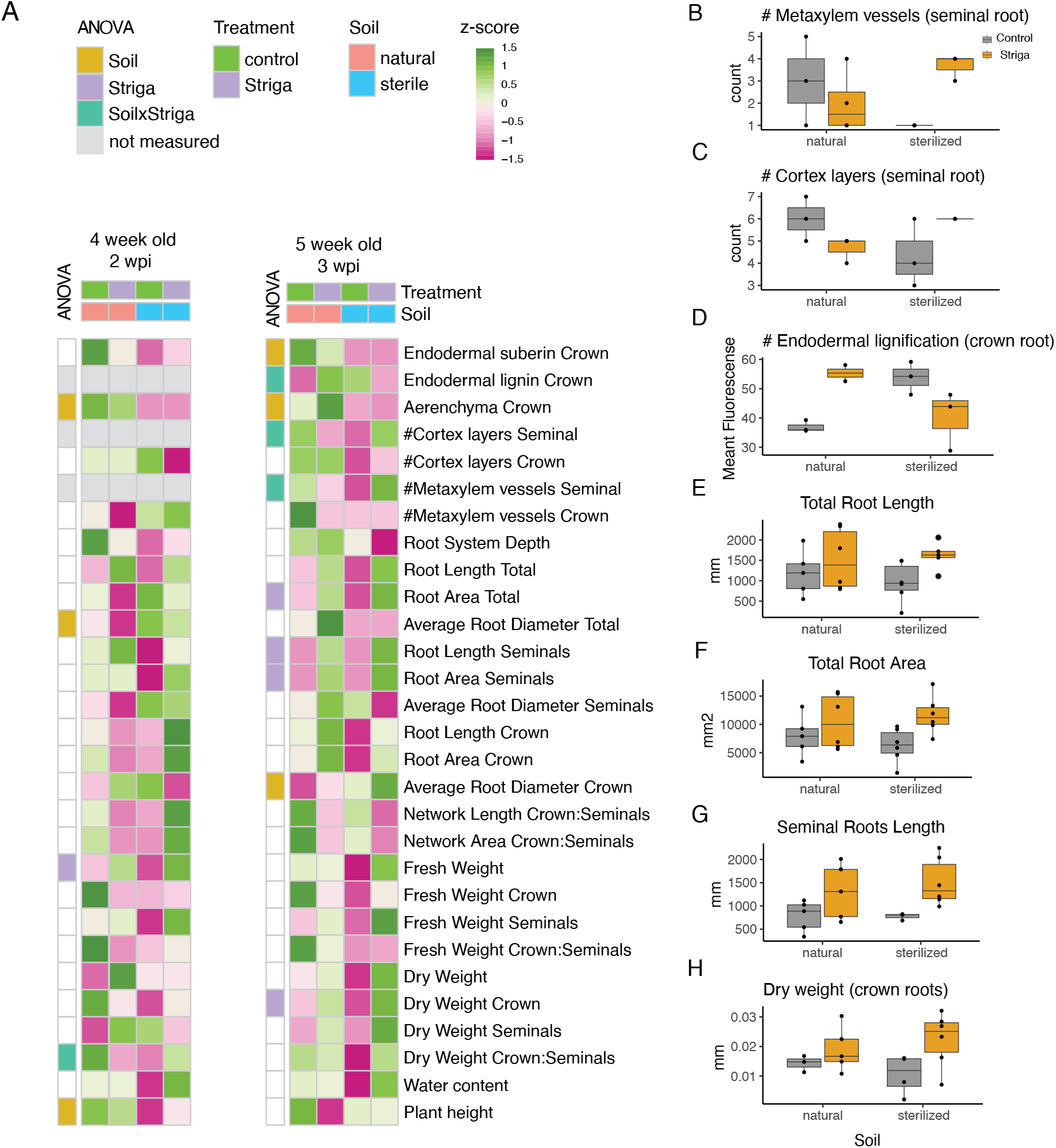
Phenotypic characterization of root cellular anatomy, root system architecture and root biomass of Shanqui Red (SQR). (A) Heatmap presents values of each trait scaled across conditions tested. The left panel of each heatmap indicates whether the trait was significantly affected by soil, Striga, and soil by Striga interaction (as identified by a two-way ANOVA). (B) The number of metaxylem vessels and (C) cortex layers in seminal roots, (D) endodermal lignification of crown roots, (E) Length and (F) area of total root system, (G) length of seminal roots and (H) dry biomass of crown roots. Data in B-H are from five-week-old plants (three weeks post-infection). The boxplots denote data spanning from the 25th to the 75th percentile and are centered to the data median. Dots represent individual values.

**Supplementary Figure 3.**
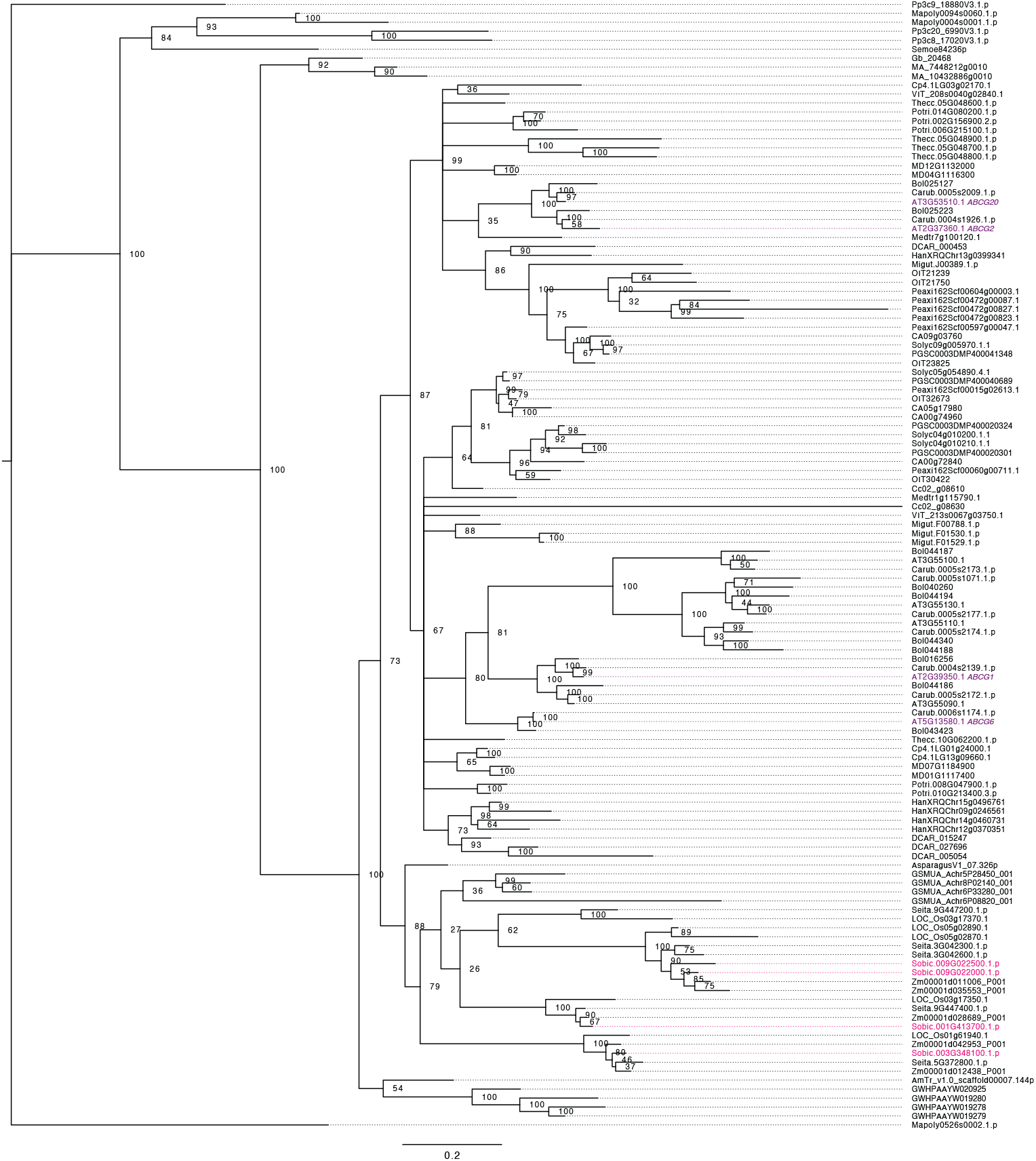
Phylogenetic tree of ABCG transporters. Phylogenetic trees generated using protein sequences of several plant species. *Arabidopsis thaliana* genes are highlighted in purple. *S. bicolor* genes are highlighted in pink for reference. AmTr: *Amborella trichopoda,* AT: *Arabidopsis thaliana,* Asparagus: *Asparagus officinalis,* Azfi: *Azolla filiculoides,* Bol: *Brassica oleracea,* Carub: *Capsella rubella,* CA: *Capsicum annuum,* Cc: *Coffea canephora,* Cp: *Cucurbita pepo,* DCAR: *Daucus carota,* Gb: *Ginkgo biloba,* HanXRQ: *Helianthus annuus,* MD: *Malus domestica,* Mapoly: *Marchantia polymorpha,* Medtr: *Medicago truncatula,* Migut: *Mimulus guttatus,* GSMUA*: Musa acuminata,* OIT: *Nicotiana attenuata,* GWHPAAYW: *Nymphaea colorata,* LOC_Os: *Oryza sativa japonica,* Peaxi: *Petunia axillaris,* Pp: *Physcomitrella patens,* MA: *Picea abies,* Potri: *Populus trichocarpa,* Semoe: *Selaginella moellendorffii,* Seita: *Setaria italica,* Solyc: *Solanum lycopersicum,* PGSC: *Solanum tuberosum,* Sobic: *Sorghum bicolor,* Thecc: *Theobroma cacao,* VIT: *Vitis vinifera,* Zm: *Zea mays.*

**Supplementary Figure 4.**
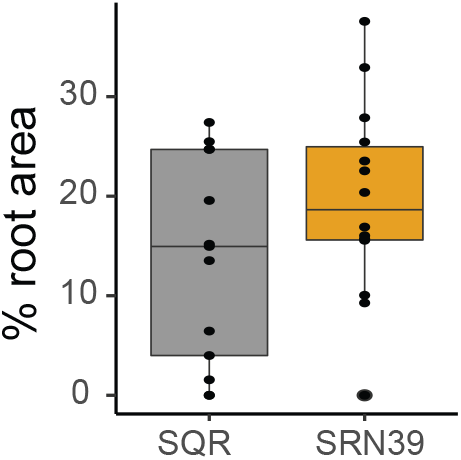
The proportion of aerenchyma relative to root cross-sectional area of 10-day-old seedlings of SQR and SRN39, measured at 7 cm from the root tip. The boxplots denote data spanning from the 25th to the 75th percentile and are centered to the data median. Dots represent individual values.

**Supplementary Figure 5.**
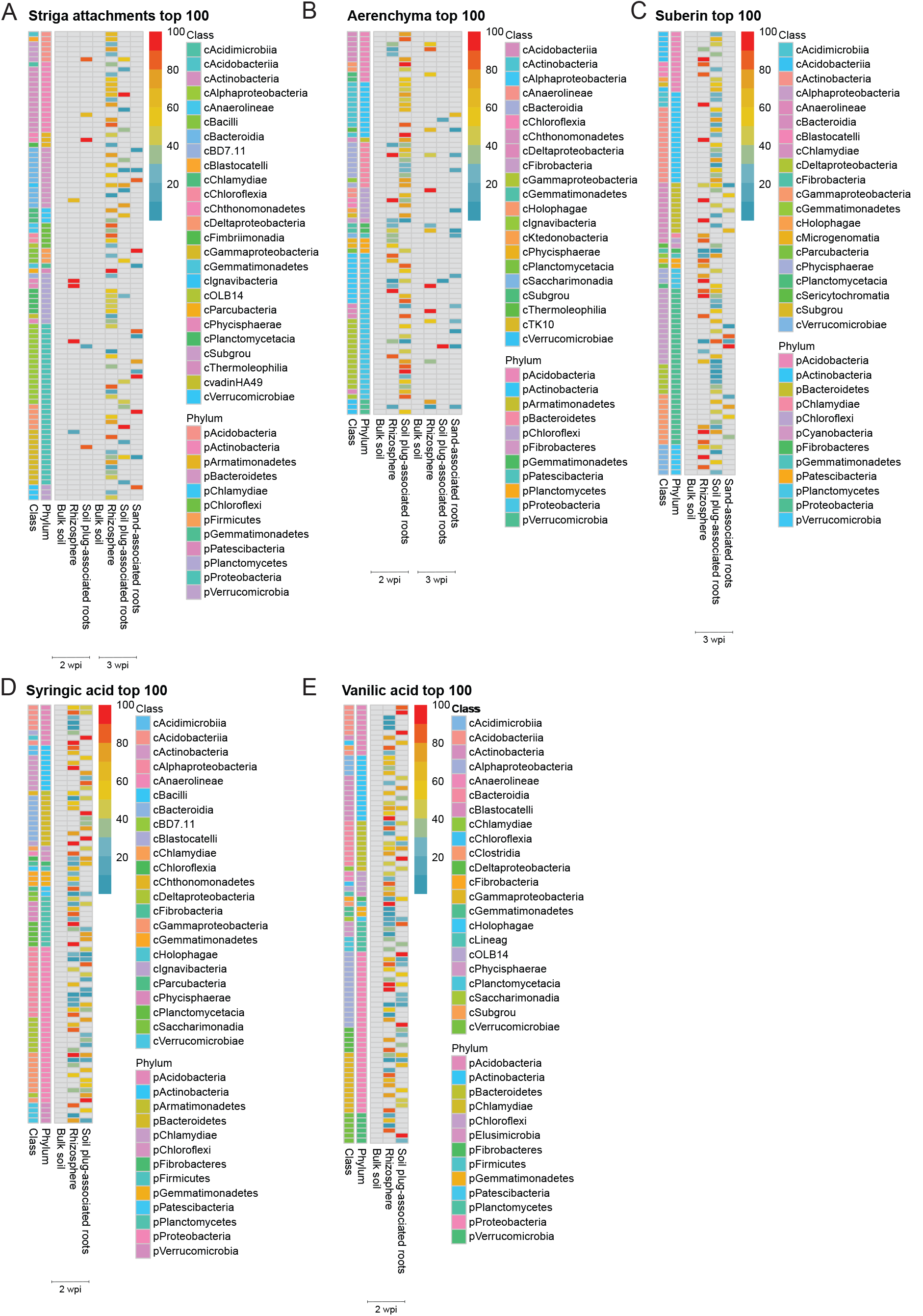
Overview of top 100 bacterial taxa predicted to (A) reduce the number of Striga attachments; (B) induce aerenchyma formation; (C) induce endodermal suberization; reduce levels of, (D) syringic acid, (E) vanillic acid. Heatmaps present the rank calculated for each taxa (with blue indicating highest, while red the lowest rank) and the microbial subcategories where each taxon was found (x-axis). Taxonomic membership at the phylum and class level is indicated with side panel colors.

**Supplementary Figure 6.**
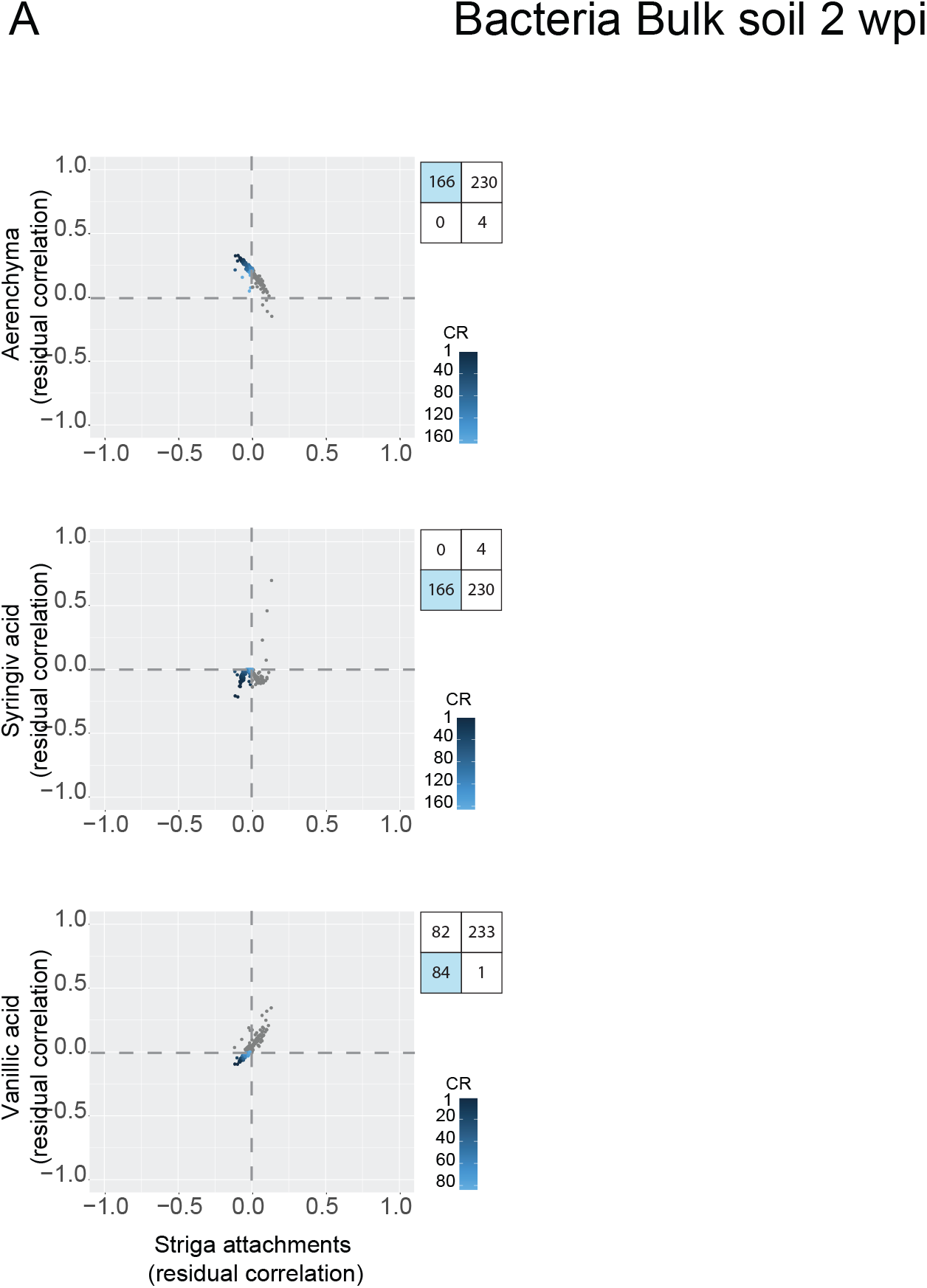
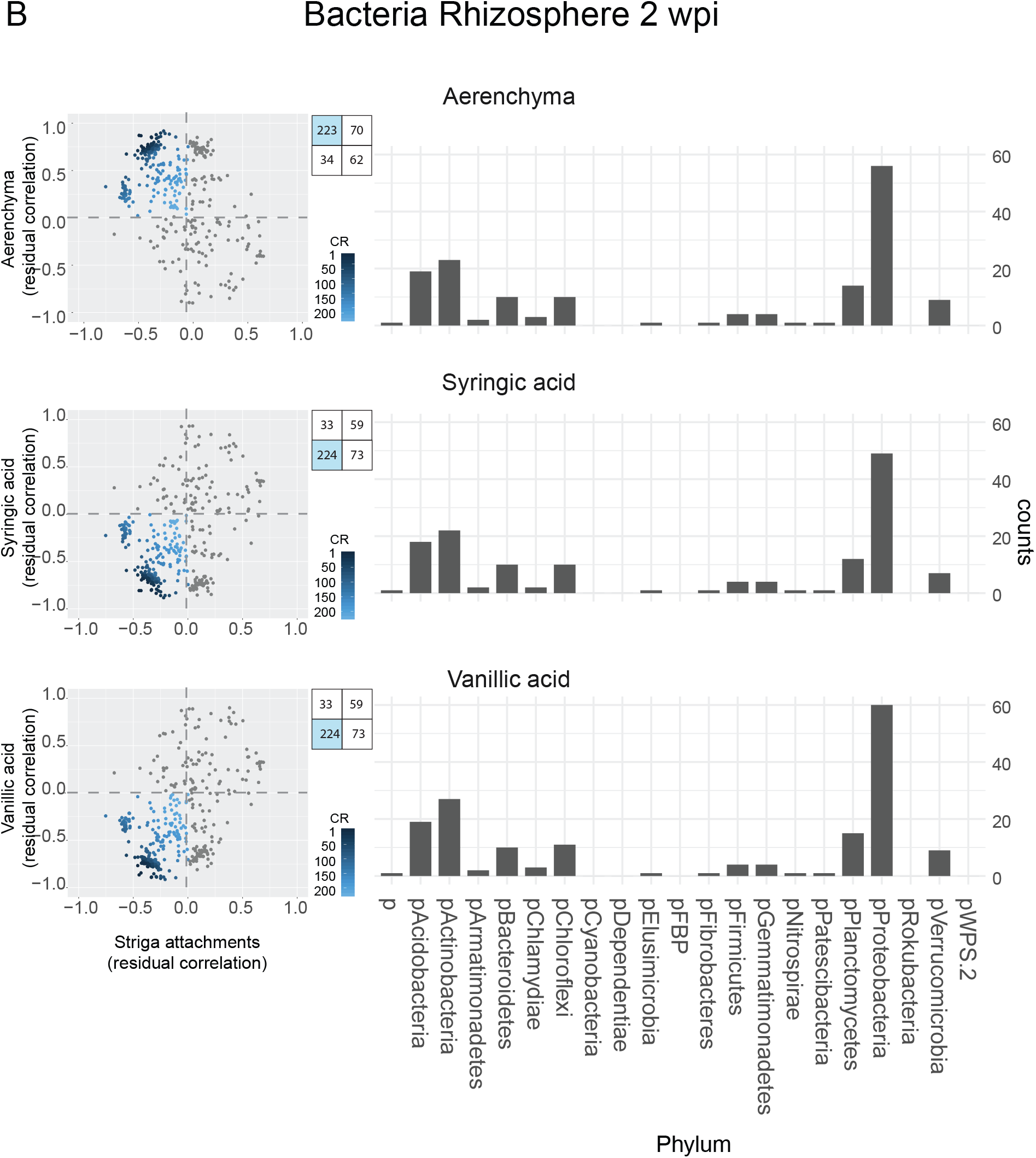
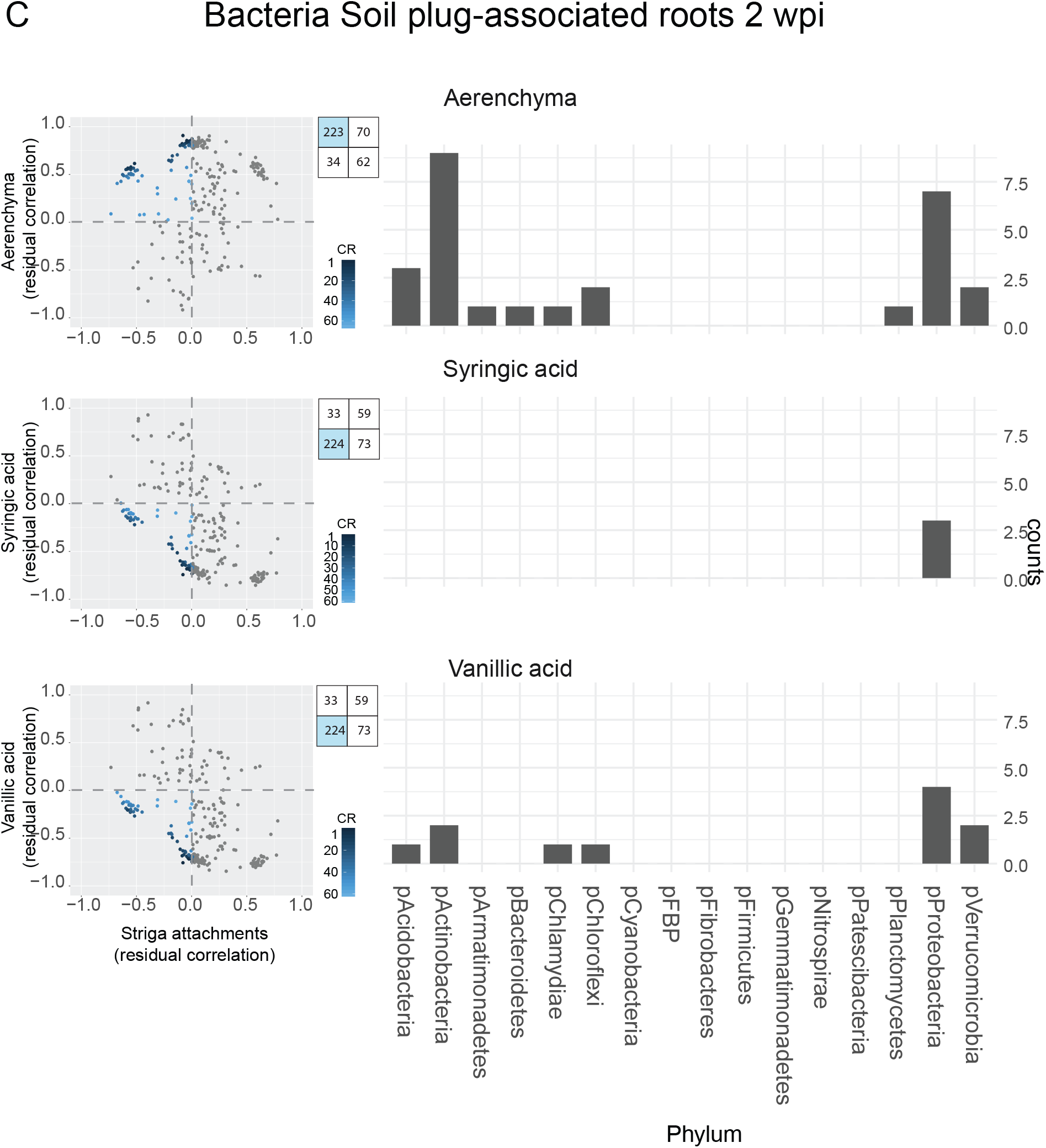
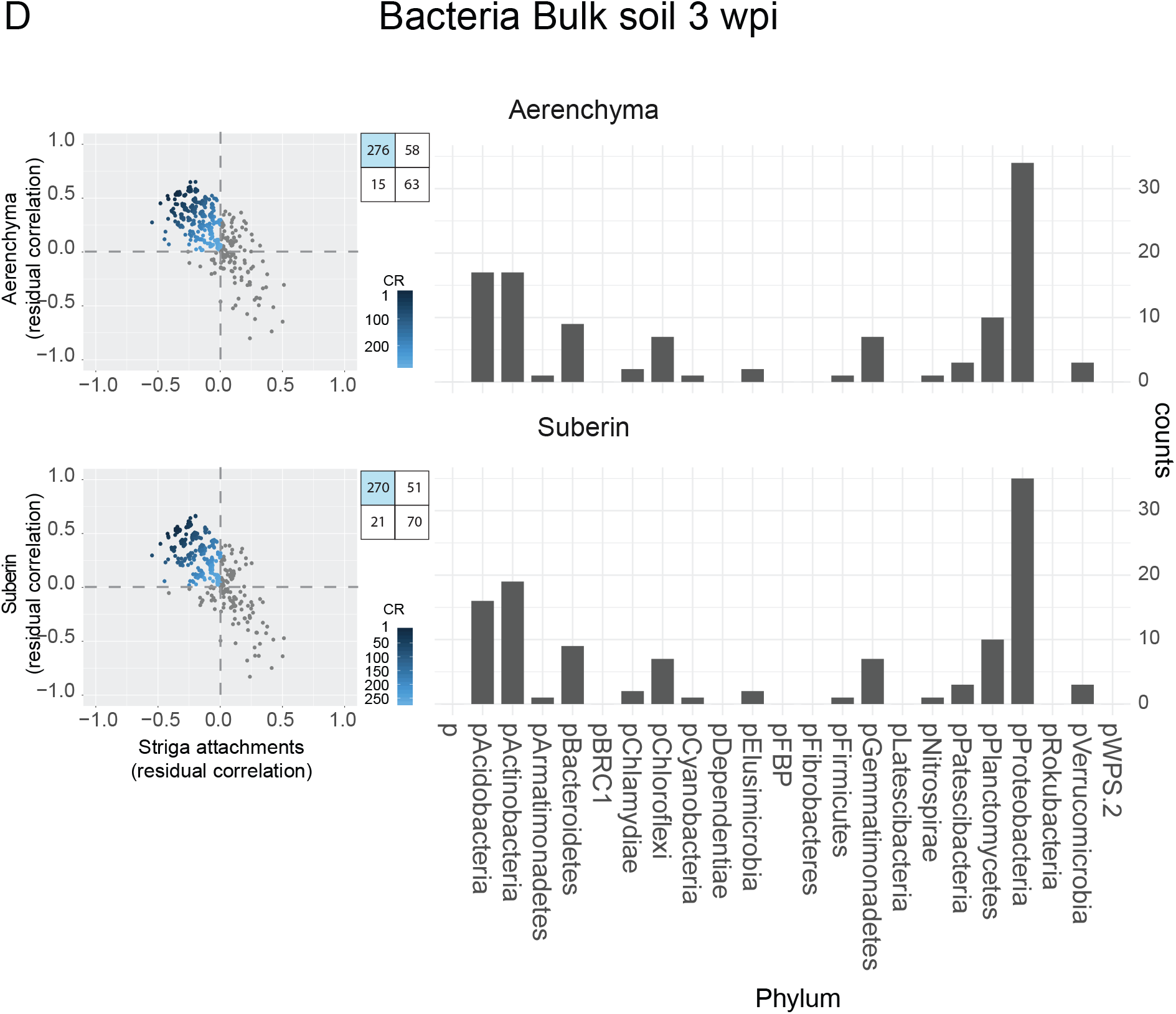
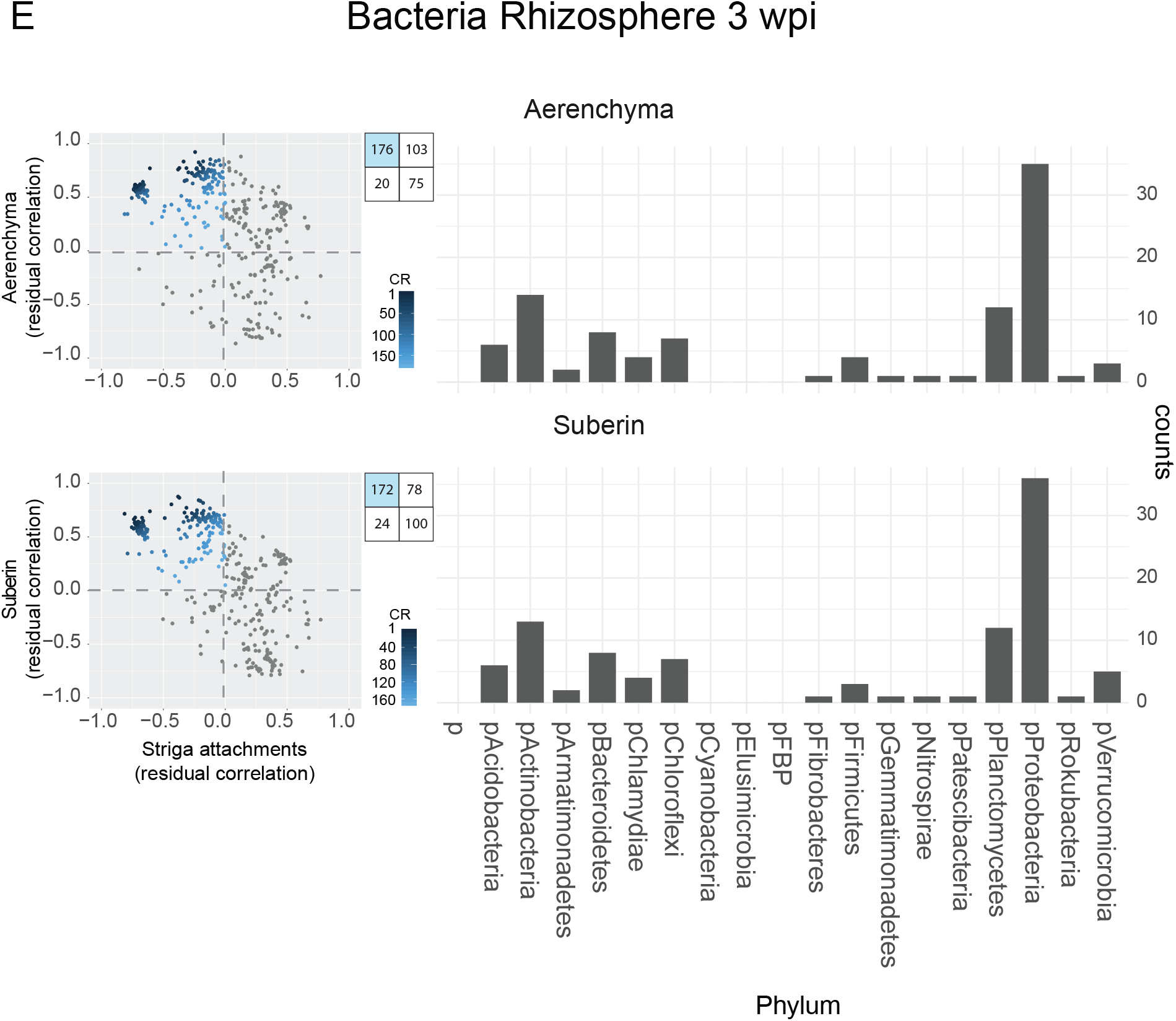
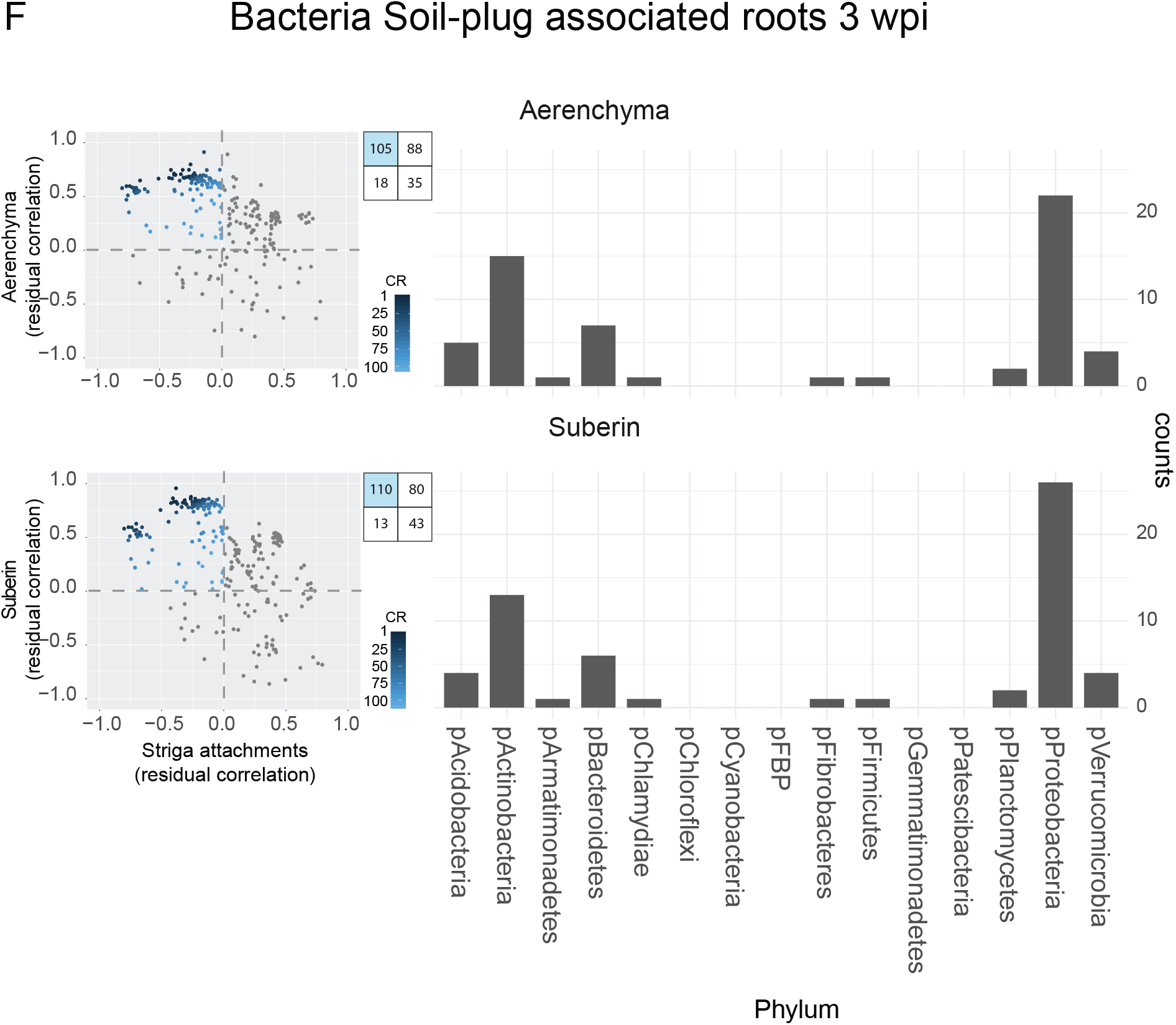
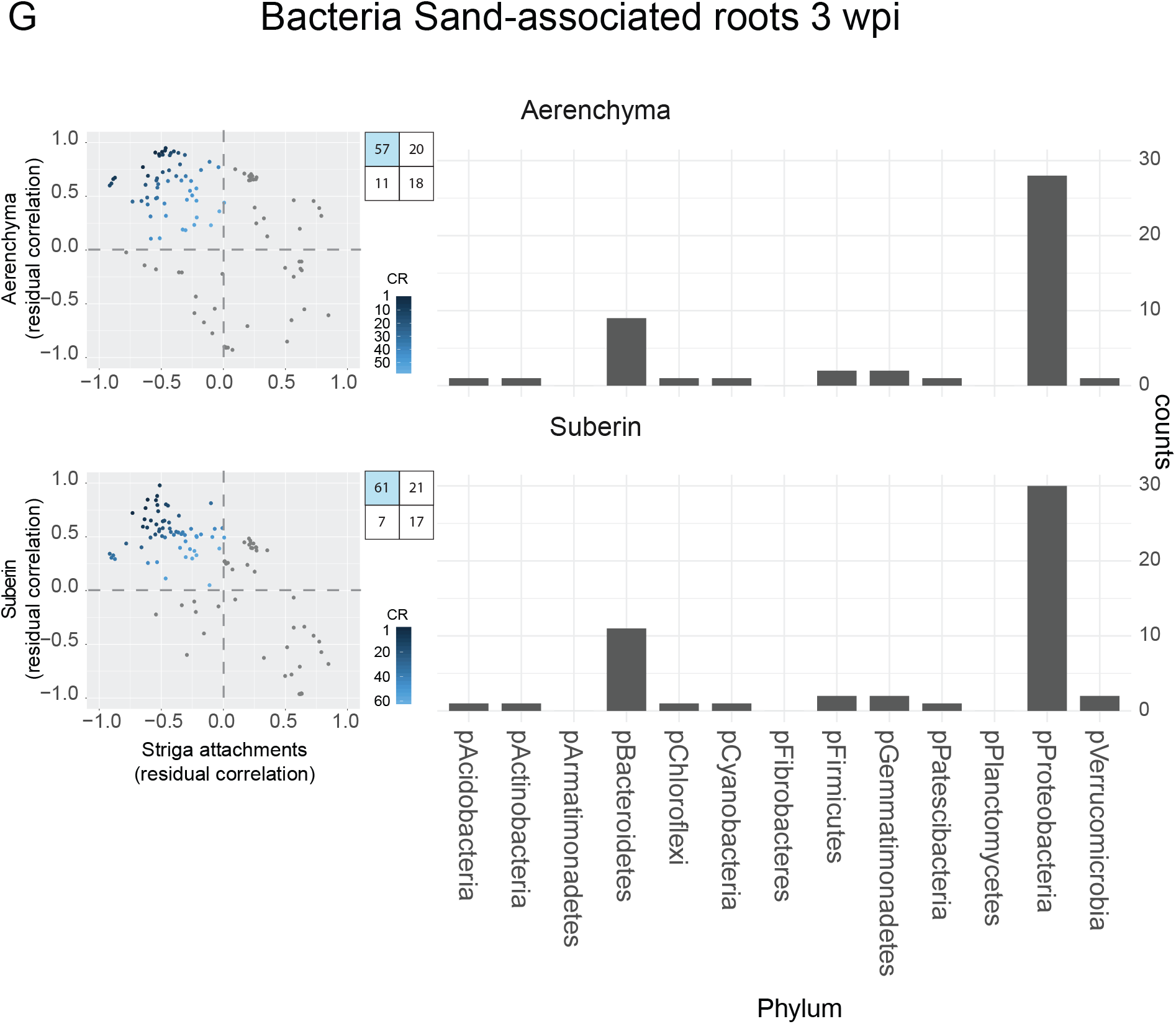
Overview of bacterial taxa found at two weeks post-infection in (A) bulk soil, (B) rhizosphere, (C) soil plug-associated roots and three weeks post-infection in (D) bulk soil, (E) rhizosphere, (F) soil plug-associated roots, (G) sand-associated roots, predicted to influence Striga infection via each of identified mechanisms (see Methods). Left panel: each dot represents individual bacterial taxon and its residual correlation found for Striga attachment (x-axis) and one of the identified mechanisms (y-axis). The four-square inset indicates number of taxa found to be: (i) negatively correlated with Striga attachment number and positively with each mechanism (left, upper square), (ii) positively correlated with Striga attachment number and positively with each mechanism (right, upper square), (iii) negatively correlated with Striga attachment number and negatively with each mechanism (left, lower square), (iv) positively correlated with Striga attachment number and negatively with each mechanism (left, upper square). The blue shading in the four-square inset indicates the number of taxa which were used for combined ranking and which represent taxa predicted to reduce Striga infection given the trait under study. Within the residual correlation plots, the intensity of blue represents their combined rank value (CR). Right panel: Number of bacteria from each phylum found to reduce Striga infection via each of the mechanisms with the cut-off of residual correlation −0.2 for Striga attachments, syringic acid and vanillic acid levels and 0.2 for aerenchyma proportion and suberin content. No bacteria passed the threshold in bulk soil 2wpi (A).

**Supplementary Figure 7.**
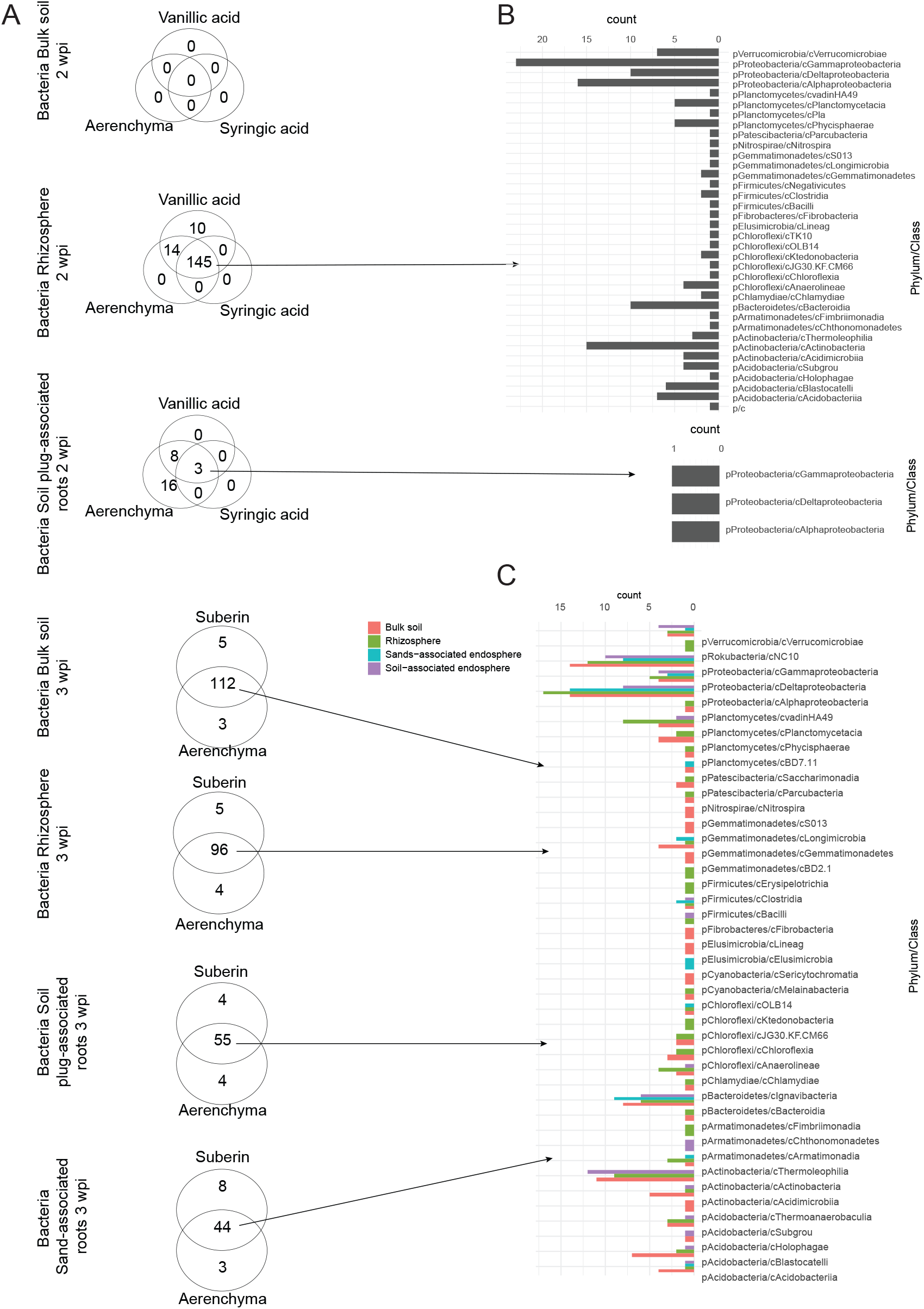
Overlap of the number of bacterial taxa predicted to reduce Striga infection via each mode of action. (A) Number of bacteria found to reduce Striga infection via each of the mechanisms with the cut-off of residual correlation −0.2 for Striga attachments, syringic acid and vanillic acid levels and 0.2 for aerenchyma proportion and suberin content. Phylogenetic membership of taxa inducing (B) all four modes of Striga suppression in the rhizosphere two weeks post-infection and inducing suberin content and aerenchyma formation across microbial sub-categories three weeks post-infection.

**Supplementary Figure 8.**
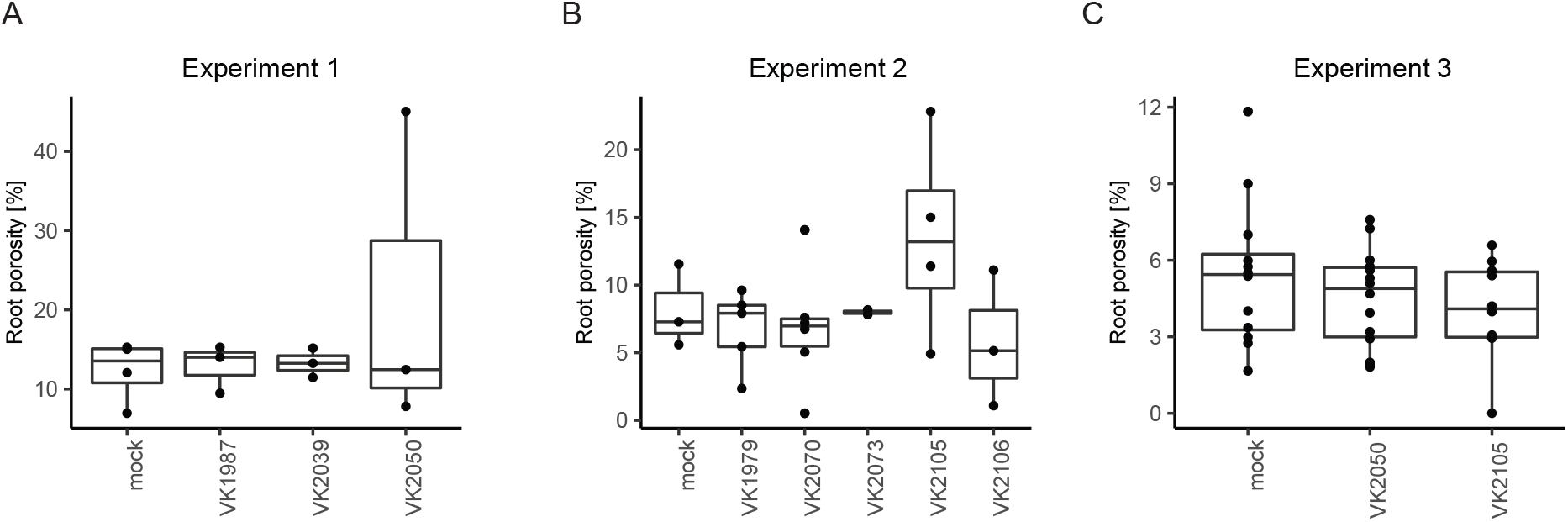
Root porosity as a proxy for aerenchyma content of the whole root system (expressed as a proportion of the volume of the whole root system) of plants inoculated with (A) Pseudomonas 1987, 2039 and 2050, (B) Arthrobacter VK1979, VK2073, VK2105, VK2105 and Pseudomonas VK2070, n = 6. (C) Isolates VK2050 and VK2105 were retested with n =15. One-way ANOVA was used to determine the effect of the inoculation. No significant differences were detected.

**Supplementary Figure 9.**
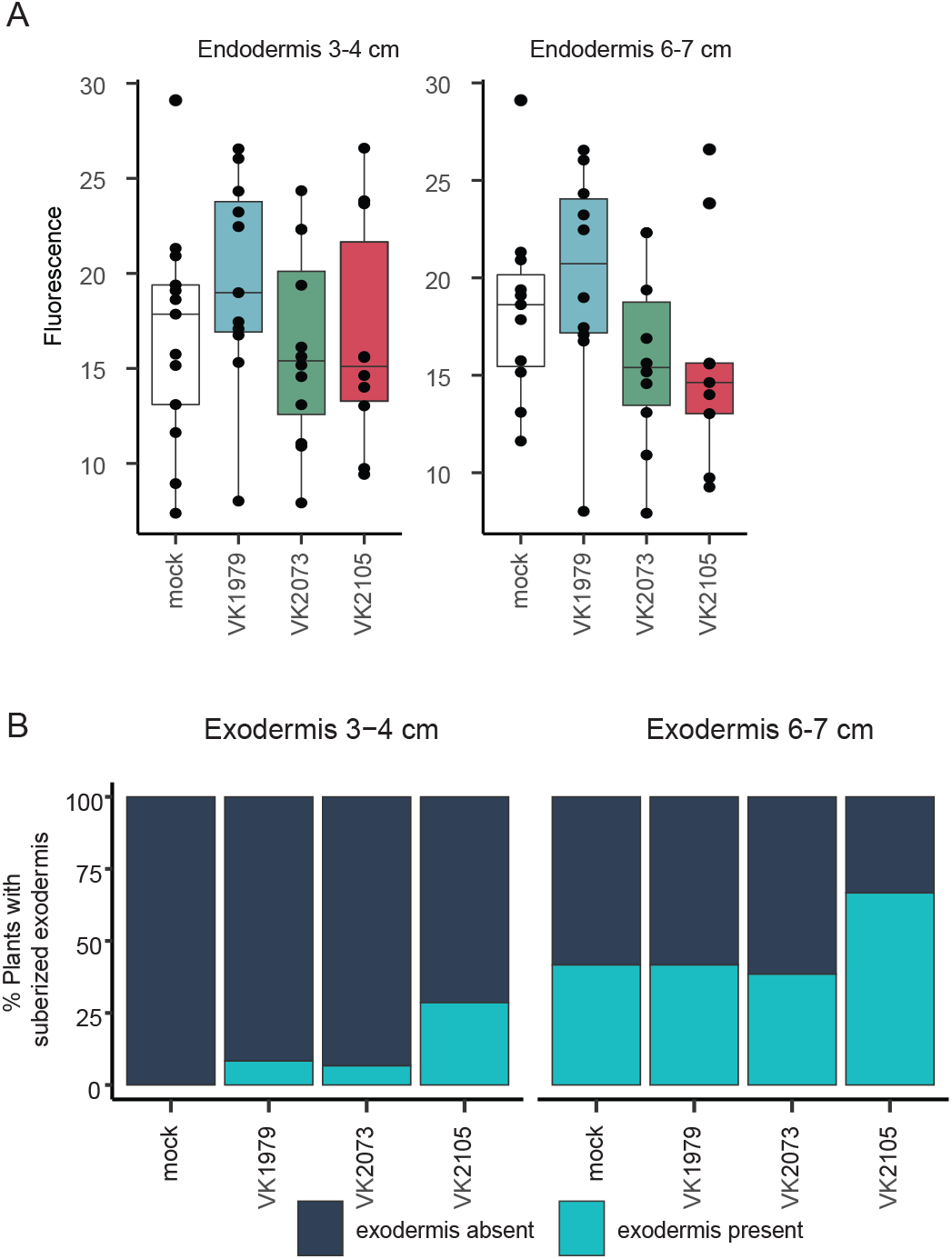
Suberin content in the main root endodermis (A) of plants inoculated with Arthrobacter strains VK1979, VK 2073 and VK 2105. Suberin was stained with fluorol yellow and quantified with mean intensity of pixel. One-way ANOVA was used to determine the effect of the inoculation. No significant differences were detected. B) Percentage of plants with a suberized or non-suberized exodermis in the root region 3-4 cm and 6-7 cm from the root tip, upon inoculation with Arthrobacter strains VK 1979, VK 2073 and VK 2105. No significant differences were detected as per Fisher exact test.

**Supplementary Figure 10.**
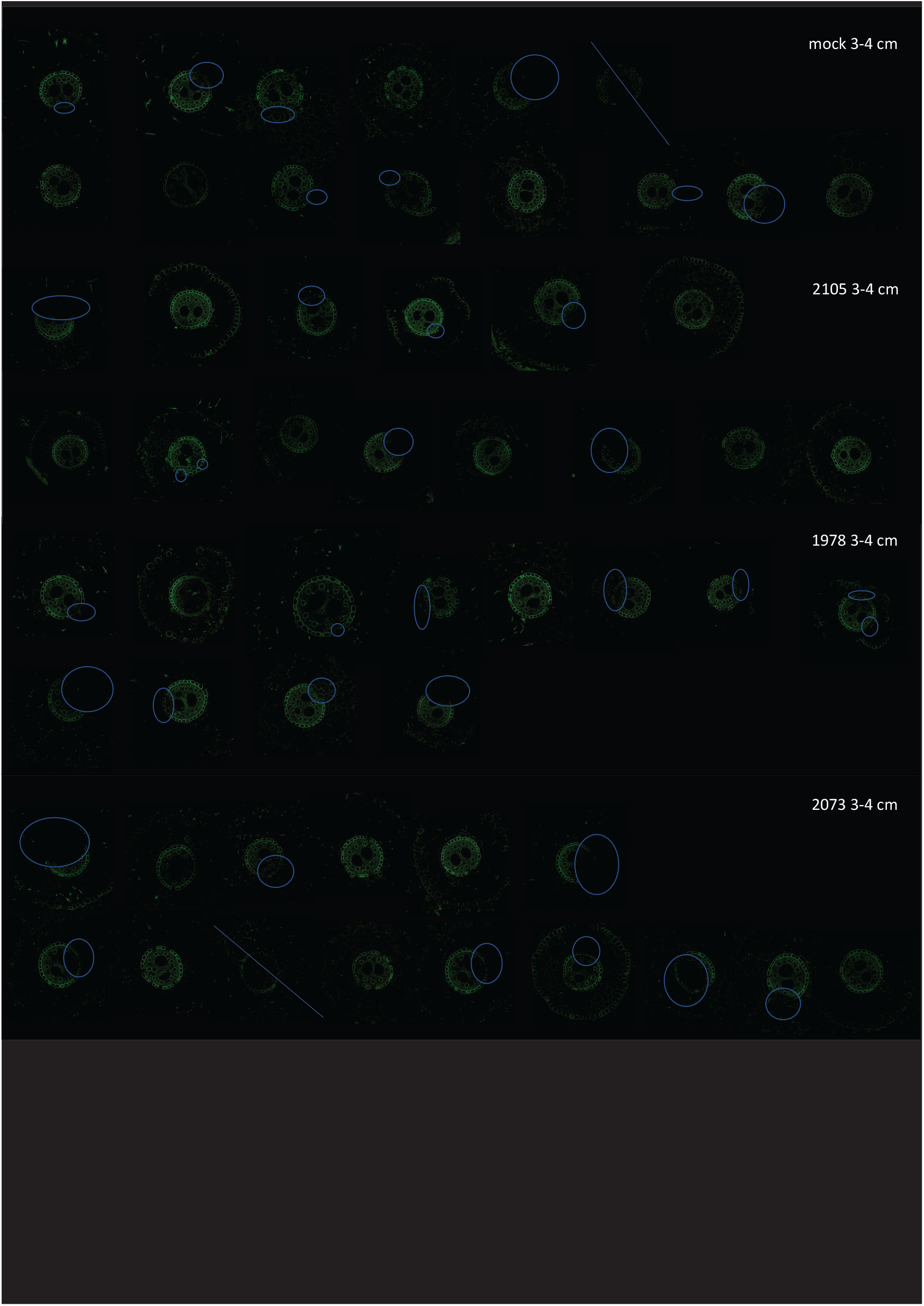
Raw images used for quantification of the proportion of suberized cells in endodermis. Blue circle denotes regions that were excluded from the analysis (see Methods).

